# A single helix repression domain is functional across eukaryotes

**DOI:** 10.1101/2022.05.09.491245

**Authors:** Alexander R. Leydon, Román Ramos Baez, Jennifer L. Nemhauser

## Abstract

The corepressor TOPLESS (TPL) and its paralogs coordinately regulate a large number of genes critical to plant development and immunity. As in many members of the larger pan-eukaryotic Tup1/TLE/Groucho corepressor family, TPL contains a Lis1 Homology domain (LisH), whose function is not well understood. We have previously found that the LisH in TPL—and specifically the N-terminal 18 amino acid alpha-helical region (TPL-H1) —can act as an autonomous repression domain. We hypothesized that homologous domains across diverse LisH-containing proteins could share the same function. To test that hypothesis, we built a library of H1s that broadly sampled the sequence and evolutionary space of LisH domains, and tested their activity in a synthetic transcriptional repression assay in *Saccharomyces cerevisiae*. Using this approach, we found that repression activity was highly conserved and likely the ancestral function of this motif. We also identified key residues that contribute to repressive function. We leveraged this new knowledge for two applications. First, we tested the role of mutations found in somatic cancers on repression function in two human LisH-containing proteins. Second, we validated function of many of our repression domains in plants, confirming that these sequences should be of use to synthetic biology applications across eukaryotes.

## INTRODUCTION

Transcriptional repression is enacted through a diverse array of mechanisms, which are often directed by a group of proteins known as corepressors (1–4). Corepressors do not directly bind DNA, but instead recruit inhibitory machinery to specific loci via interactions with transcription factors. Among the best-studied corepressors are: animal SMRT (silencing mediator of retinoic acid and thyroid hormone receptor) and NCoR (nuclear receptor corepressor) complexes (5, 6); yeast Tup1 (7–9) and its homologs *Drosophila* Groucho (Gro) and mammalian transducing-like enhancer (TLE) (10); plant TOPLESS/TOPLESS-RELATED1-4 (TPL/TPR1-4), LEUNIG/ LEUNIG_HOMOLOG (LUG/LUH), and HIGH EXPRESSION OF OSMOTICALLY RESPONSIVE GENES 15 (HOS15) (11–16). Despite knowing the identity of many corepressor proteins, much is left to uncover about how these complexes integrate input signals to create, sustain, and relieve transcriptional repression.

Plant corepressor families share a general structural similarity, where the N-terminus contains protein-protein interaction domains followed by an unstructured linker to a C-terminal WD40 beta-propeller domain (13, 14). While the full N-terminal structure is not highly conserved, all plant corepressors share a Lis1 Homology domain (LisH), which is generally known as a protein multimerization interface (17–25). In TPL, the LisH enables homodimerization of TPL/TPRs (26, 27). The LisH is followed by a <u>C</u>-terminal to <u>L</u>is<u>H</u> (CTLH) domain that binds partner proteins via an ETHYLENE-RESPONSIVE ELEMENT BINDING FACTOR-ASSOCIATED AMPHIPHILIC REPRESSION (EAR) motif (12, 28). The N-terminal domain of TPL also contains a CT11-RanBPM (CRA) domain, which acts as a second homo-multimerization interface that folds back over to stabilize the LisH domain (26, 27). In previous work, we found that the N-terminal domain of TPL contains two distinct repression domains, one of which is the LisH domain (29). Specifically, the first of two alpha helical regions within the LisH, termed hereafter Helix 1 (TPL-H1), was sufficient to repress transcription in yeast (29).

The 33-residue LisH motif is found in many proteins across eukaryotes; currently, there are more than 25,000 unique LisH sequence entries in the SMART protein database (30). These proteins have a broad range of annotated functions, including: cytoskeleton-interacting proteins, ubiquitin ligase complexes, and transcriptional regulation. The founding member LIS1 regulates microtubule function and is required for proper neurodevelopment (31). While LIS1 has been broadly studied in its cytoplasmic context, recent work has also demonstrated a nuclear role in gene expression (32). Several E3 ubiquitin-ligase-associated proteins carry LisH domains, such as DDB1–Cul4-associated factor 1 (DCAF1, (33)), which is involved in myriad pathways contributing to development and disease (34). The glucose-induced-degradation (GID) E3 ligase complex is assembled by intermolecular LisH interactions (24). Other LisH-containing proteins have well characterized roles in human health and disease such as the oncogene Transducin-beta like 1 (35). TBL1 is a component of the SMRT/NCoR complex (6), and acts as an exchange factor, facilitating the conversion of SMRT/NCoR repressed loci into sites of active transcription (36). TBL1’s LisH domain is required for its transcriptional activity (22). Understanding conserved functions of LisH domains has the potential to simultaneously shed light on multiple core cell and developmental processes and provide insights into diseases that result from dysregulation of LisH-containing proteins.

Previously, we have recapitulated the auxin response pathway in yeast by transferring essential components from plants (*<u>A</u>rabidopsis thaliana* <u>A</u>uxin <u>R</u>esponse <u>C</u>ircuit in *Saccharomyces cerevisiae, At*ARC^*Sc*^; (37)). In *At*ARC^*Sc*^, an auxin-responsive transcription factor (ARF) binds to a promoter driving expression of a fluorescent reporter. ARF activity is repressed by interaction with a fusion protein comprised of an Aux/IAA protein and the N-terminal 100 amino acids of TPL. Reporter activation can be quantified after addition of auxin by flow cytometry (37). In this way it is possible to test the direct effect of various mutations in TPL, or other transcriptional repressors, at an orthogonal, synthetic locus in a quantitative manner.

Here, we modified the *At*ARC^*Sc*^ to better understand the repressive function and evolutionary history of LisH-H1s. We first interrogated the critical residues within AtTPL-H1 that are required for robust transcriptional repression. Next, we built a library of H1 sequences from proteins with diverse annotated functions across the eukaryotic lineage to test the extent of H1 repressive function. We then focused our attention on two applications. We used our yeast assay to survey the effect of documented somatic cancer variants of two human LisH-containing proteins (HsTBL1 and HsDCAF1) on protein stability and repressive function, and we tested our most repressive H1 sequences as candidates for synthetic transcriptional repressors in plants. Together, our findings uncovered the ancestral transcriptional repression ability of the LisH domain, and showcased how this system can be used to understand disease states, as well as be incorporated into synthetic biology applications.

## RESULTS

### The TPL LisH domain is a short transcriptional repression domain

By directly fusing corepressor fragments to an auxin co-receptor (IAA3 in these studies) and thereby bringing it in close proximity to the synthetic auxin-regulated locus, the *At*ARC^Sc^ makes it possible to sensitively measure repressive activity even in fragments that would not be recruited to DNA on their own. Here, we focused on the small modular LisH domain which we previously demonstrated is sufficient to repress transcription in *At*ARC^Sc^ assays (29, 37), even when truncated to its first 18 amino acids (H1, Figure 1A-B). We first identified solvent-facing amino acids in AtTPL-H1, as these residues were less likely to be involved in stabilizing the hydrophobic interactions between intra-AtTPL helical domains and might be available to interact with partner proteins (26, 27). Six of these candidate residues were mutated to alanine (Fig. 1A, pink residues) in the context of H1-IAA3, and assayed for repression activity. The amino acids on either end of the helix (R6 and E18) were required for repression (Fig. 1B), as we observe high reporter expression when these are replaced with alanines. A mutation of E7, the immediate neighbor of R6, slightly increased reporter expression (Fig. 1B) and lowered the final activation level after auxin addition when compared with wild type AtTPL-H1. This result suggests it likely plays a smaller, supporting role in repressive function (Fig. 1C). Consistent with this interpretation, the R6A, E7A double mutant behaved similarly to R6A alone. Likewise, the D17A, E18A double mutation did not enhance the effect of E18A alone. Q14A and D17A were indistinguishable from wild type H1 (Fig. 1C). In contrast to the other mutations which had negative if any effects on H1 repression activity, F10A strengthened the durability of repression H1, converting it into an auxin-insensitive repression domain (Fig. 1C). One explanation for this stabilizing effect is that in the context of the full-length AtTPL protein F10 is buried in a hydrophobic cluster (along with F163, F33, F34 and L165, (26, 27)). Truncations of AtTPL left F10 solvent-exposed, rendering it less stable overall. This situation was at least partially remedied by substituting a smaller, less hydrophobic residue (F10A).

**Figure 1.**
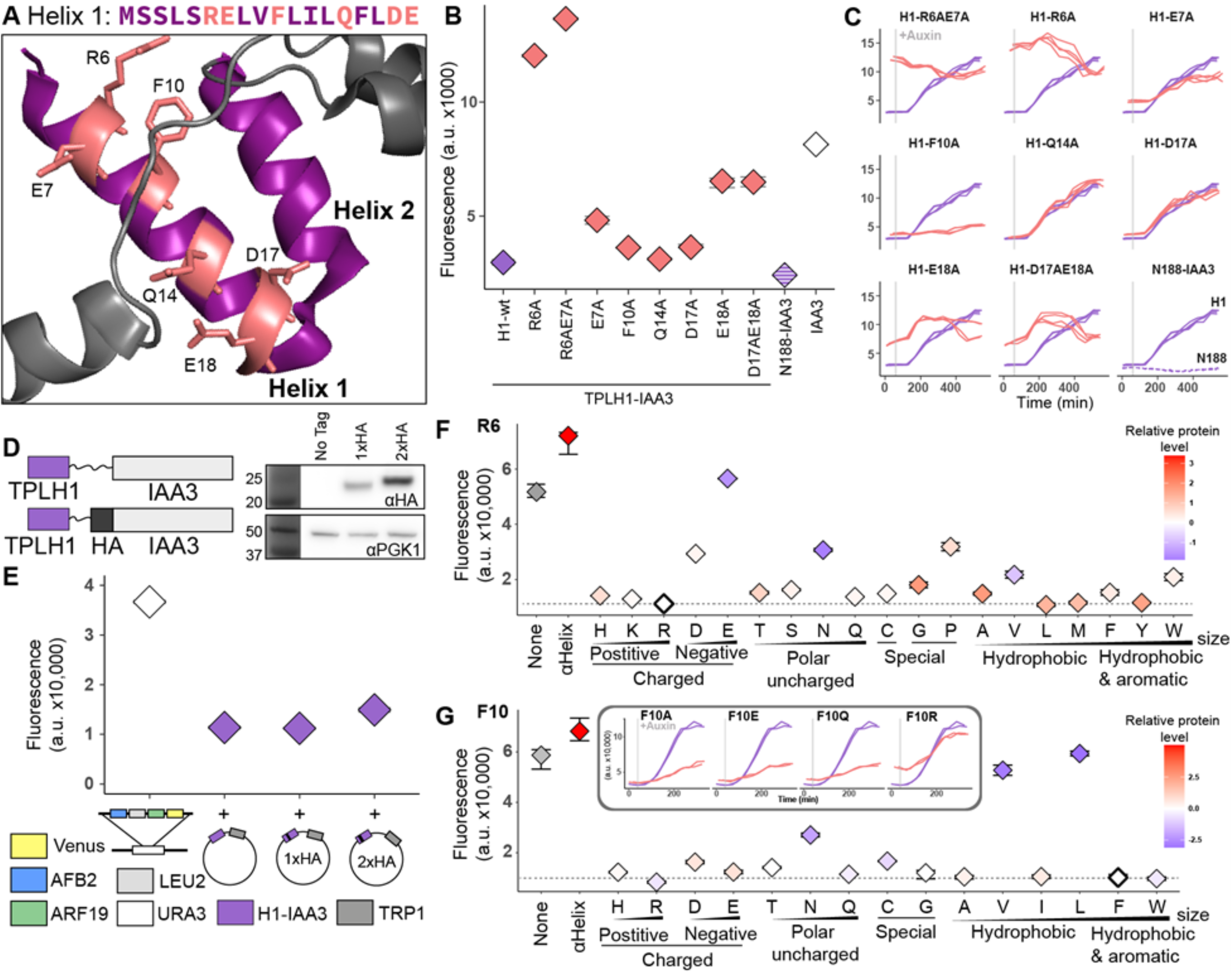
AtTPL LisH H1 is a very short autonomous repression domain. **A**. Sequence and structure of Helix 1 (AtTPL-H1) (PDB: 5NQS). The LisH domain is colored purple, and amino acids chosen for mutation are highlighted in both the sequence and the structure with pink. **B**. An alanine scan of residues predicted to be solvent-facing. Repression activity of indicated alanine substitutions is in red, and wild-type H1 sequence is in blue. N188-IAA3 (blue hatch) and IAA3 with no corepressor (white) are included for reference. **C**. Time course flow cytometry of selected H1 mutations. Auxin (10µM IAA) was added at the time indicated by the gray bar. **D**. Schematic of HA epitope placement and western blots of tagged constructs. **E**. Repression activity of HA-tagged AtTPL-H1 constructs. All components of the *At*ARC^Sc^ that were held constant across experiments (*auxin promoter::Venus, ARF19, AFB2*) were integrated at the URA3 locus. The variable H1-HA-IAA3 constructs were expressed from a plasmid carrying the *TRP1* prototrophic gene. **F-G**. Flow cytometry was used to measure the repression activity of AtTPL-H1 constructs with a range of amino acid substitutions at position R6 (**F**) and F10 (**G**), as alanine substitutions at these positions strongly inhibited and enhanced repression activity, respectively. Constructs with wild-type residues are indicated with a bold outline, and the dotted lines represent their repression strength. Protein accumulation was assayed by western blot using an α-HA antibody and normalized to yeast PGK1. Levels of AtTPL-H1 mutants are shown relative to wild type AtTPL-H1 with each data point color coded from blue (low) to red (high) expression on a log2 scale. **G**. [Inset] Specified variants were tested in the more sensitive, fully integrated *At*ARC^Sc^. The blue line in each graph is the wild-type TPL-H1 sequence. **B**,**C**,**E**,**F**,**G**. Each panel represents two independent time course flow cytometry experiments of the AtTPL-H1 constructs indicated, all fused to IAA3. For all cytometry, every point represents the average fluorescence of at least 10,000 individually measured yeast cells (a.u. - arbitrary units).

To rapidly test an expanded library of mutations in LisH H1 sequences, we designed a new auxin response circuit (ARC) that includes an epitope tag to allow quantification of repressor protein levels (Fig. 1D). All parts of the circuit that are held constant across experiments were integrated at the *URA3* locus. The H1-1xHA-IAA3 fusion protein was both detectable in western blots and showed minimal interference with repression activity (Fig. 1E, Supplement 1). Our next step was to perform an amino acid swap at AtTPL-H1 residues R6 and F10 (Fig. 1B-C). In the case of R6, we observed broad tolerance of amino acid swap (Fig. 1G), with the exceptions of charge inversion (R6D and R6E), and an unsurprising reduction of repression in R6P, which is likely to interfere with its alpha-helical structure. The unexpected result that R6A has only a mild decrease in repression in this experimental design is likely due to higher relative expression of a plasmid-based AtTPL-H1 compared to the other components. In the context of this altered stoichiometry of circuit components, repression strength is slightly less sensitive to loss of function in AtTPL-H1. Several amino acid swaps had a negative effect on protein accumulation (R6E, R6N, and R6V), likely explaining the observed loss of repression. In all experiments we compared these mutations to a well characterized alpha-helical linker sequence (□-helix-HA-IAA3) as a control, as well as IAA3 alone (None) (Fig. 1F,G).

In the case of F10, we also observed a broad tolerance of amino acid substitutions (Fig. 1G). A small number of substitutions had negative effects on protein accumulation (F10N, F10V, and F10L) which likely explained their reduced repressive activity. Several substitutions (E, Q, and R) showed similar repression strength to wild type, despite their different physicochemical character. We introduced these variants into the fully integrated *At*ARC^Sc^ to test for sensitivity to auxin treatment. Similarly to F10A, substitution of a negative (E) or polar (Q) residue in TPL-H1 resulted in an increased durability of repression; in contrast, the substitution of a positive charge (R) had a milder effect (Fig. 1G inset). These results support the hypothesis that the exposure of a hydrophobic residue (F) by truncation negatively impacts TPL-H1 activity. In addition, applications using the TPL-H1 as an autonomous repression domain should incorporate one of the stronger repression variants assayed here.

### Defining the LisH H1 sequence

Although the repressive function of most LisH-H1 domains have not been directly tested, many LisH-containing proteins are known transcriptional regulators (16, 38–41). Many other LisH-containing proteins are primarily cytoplasmic, or have well-studied primary functions in ubiquitination or cytoskeletal dynamics, making it unlikely anyone would have tested their impact on transcription. Recent work on Lis1, which contains the founding LisH domain and has been thought to be exclusively cytoplasmic, suggests that this assumption about functions for these proteins may be misplaced. New evidence suggests that Lis1 moonlights as a transcriptional repressor (32), raising the possibility that the same might be true for other LisH-containing proteins.

To better understand the diversity of LisH-H1 sequences and the proteins that carry them, we performed Maximum Likelihood (ML) reconstruction (42) of sequences sampled from diverse LisH-containing proteins across eukaryotes (Fig. 2A, Supplement. 2). While the 18 amino acids of the LisH-H1 sequences is too short to produce optimal bootstrap values (43), our reconstruction allows us to make associations based on sequence similarity. The sequences cluster into five main clades, defined by nodes I-V. As some LisH-H1 sequences were identical across orthologous genes, a given label for a LisH sequence may be only one of many possible labels (Supplement 3). To further check for functional sequence associations, we annotated the listed protein’s functions and cellular localization (Fig. 2B). We observed a related cluster of genes with published roles in transcriptional activation (Fig. 2B, blue circles), as well as a related cluster of H1 sequences from published repressors (Fig. 2B, red circles). Sequence alignments highlight residues of interest across genes and clades (Fig. 2C). For example, there are several highly conserved residues (L8, N9, L11, I12, L16, and Y15) that comprise the inward-facing dimerization interface in known structures.

**Figure 2.**
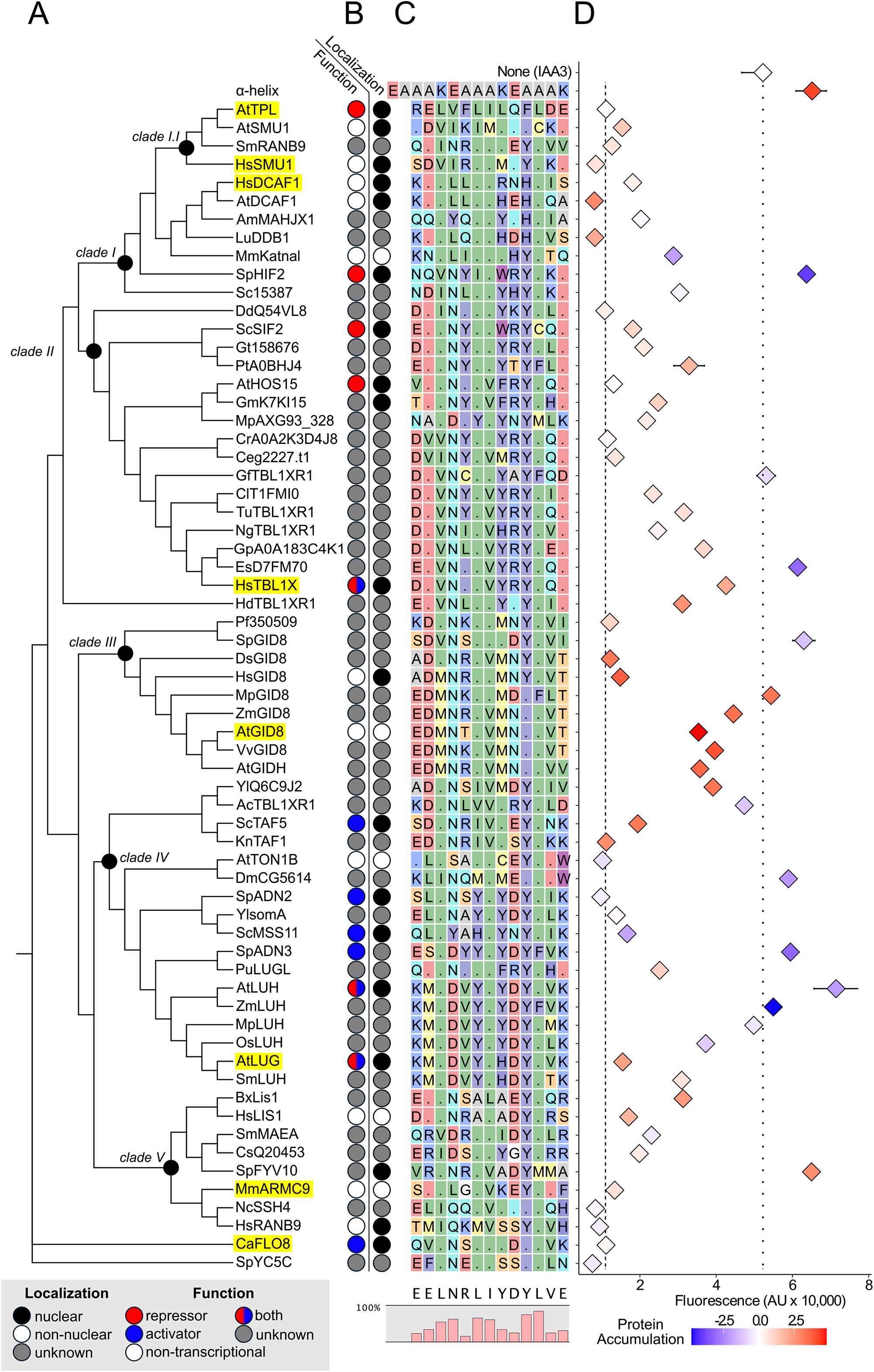
The repressive function of the LisH domain is likely ancestral. **A**. A Maximum Likelihood (46) phylogeny of LisH-H1 sequences from diverse eukaryotes. Ancestral sequences of interest were inferred (42) at nodes of interest (black dot). **B**. Published function and subcellular localization for each protein. The first column marks whether a protein is a transcriptional repressor (red), transcriptional activator (blue), has another function (white), or an uncharacterized function (gray). The second column marks proteins as nuclearly localized (black), non-nuclear (white), or uncharacterized (gray). **C**. LisH-H1 sequences were aligned and residues colored by their physicochemical class (RASMOL color scheme (49)). Residues that are the same as those in the AtTPL-H1 sequence at the top of the alignment are indicated with a period. The consensus sequence for H1, and the relative conservation rate of different residues along the helix, are displayed below the alignment. **D**. Flow cytometry and a modified *At*ARC^Sc^ depicted in Fig. 1E was used to quantify the relative repressive function of different LisH-H1s. We have marked the fluorescence levels detected by the positive AtTPL-H1-HA-IAA3 control (dashed line) and negative IAA3 repression control (dotted line). Protein accumulation was measured by western blot and normalized to yeast PGK1. Levels of protein expression are shown relative to AtTPL-H1 with each data point color coded from blue (low) to red (high) expression on a log2 scale.

### LisH repressive function appears to be widely conserved and ancestral

The modifications to *At*ARC^SC^ (Fig. 1E) allowed us to directly compare repressive activity across distantly related LisH-H1 sequences. To do so, we created a plasmid library using representative LisH-H1 sequences from across our reconstruction (Fig. 2A). We also tested protein levels for all fusion proteins by western blot. One not very surprising trend is that proteins that are detected at lower levels are generally poor at repression; however, the converse is not true: high accumulation of a fusion protein was not well-correlated with strong repression (Fig. 2D, Supplement 4).

While Clade I had high sequence diversity (Supplement 5), the overall similarity to TPL-H1 led us to hypothesize that these domains would also have a repressive function. This is indeed what we observed (Fig. 2D). Clade I LisH-H1 sequences from proteins involved in protein ubiquitination SmRANB9, DCAF1 (from *Homo sapiens* and *Arabidopsis thaliana*), and LuDDB1 (44) had repressive function, as did the splicing factor HsSMU1 (45). In addition, we found that H1s that belong to genes characterized as nuclear-localized transcriptional repressors, such as ScSIF2, AtHOS15, and AtLUG robustly repressed reporter activity to similar levels as AtTPL (Fig. 2D). However, many of the strongest repressors were found in genes across the tree without previously characterized roles in transcriptional repression (e.g., HsSMU1, SpADN2, and NcSSH4).

We observed that Clade II was characterized by a slightly lower sequence variability than Clade I and a high incidence of R14 and E18 residues (Fig. 2C, Supplement 5). LisH-H1s from the repressor proteins ScSIF2 (46) and AtHOS15 (40) repress strongly, while H1s closer in sequence to HsTBL1X are somewhat weaker. HsTBL1 recruits the repressive SMRT/NCoR complex, but upon stimulus facilitates the recruitment of transcriptional activators to the locus (36). Perhaps HsTBL1-H1, and closely related domains, have retained weaker repressor function to permit better exchange activity.

Clade III LisH-H1 sequences encompassed all predicted members of the multiprotein E3 ligase GID complex (24), also known as the CTLH (carboxy-terminal to LisH) complex in mammals (47). Clade III sequences were well conserved, with a high conservation of M13, and N14 (Fig. 2C, Supplement 5), which are on the solvent facing side, suggesting that these residues might lower affinity with repressive partner proteins. Most clade III LisH-H1s were expressed at levels higher than AtTPL, and sequences with both E6 and D7 showed the lowest capacity to repress.

Clade IV H1s included the nuclear localized transcriptional activators SpADN2, SpADN3 and ScMSS11, as well as the plant corepressors AtLUG and AtLUH (Fig. 2B). This clade was marked by a high incidence of Y11, Y13, and K18 residues and a strikingly low level of protein accumulation (Fig. 2C, Supplement 5). The L13Y variation may contribute to this loss, as L13 is a highly conserved residue elsewhere. Despite the low level of SpADN2-H1 and ScMSS11-H1, they were highly repressive.

Finally, Clade V included highly diverse sequences belonging to genes coding for both nuclear and non-nuclear localized proteins with no annotated transcriptional functions (Fig. 2B). The HsLIS1-H1 sequence is both well expressed and repressive, pointing to a possible role for this sequence in mediating repression, consistent with other recent findings (32). Interestingly, unlike ScGID8, H1s from other GID/CTLH complex members such as SmMAEA and HsRANB9 retain repressive ability (48).

To further investigate these trends, we reconstructed predicted ancestral LisH-H1 sequences for nodes of interest across the phylogeny (Fig. 2 – Supplement 6). All reconstructed sequences were good repressors except for the Clade III sequence which was poorly expressed. The Clade III reconstruction also contained the D7, M13 and N14 variations found in many poor repressors within this group.

It is striking how few residues within the LisH-H1 sequence are required for repressive function. Repressive function is found in LisH-H1s across all clades measured, as well as basal LisH-H1s such as CaFLO8-H1 and SpYC5C-H1. Residues outside of the conserved core LisH-H1 motif (comprised of the hydrophobic amino acids L8,11,16 and I12) serve mainly to tune the repressive function. This indicates that the most important determinant of repression is the multimerization interface and suggests that most LisH-H1 sequences should retain this activity if localized to chromatin. It is worth noting that in these assays we are only testing the LisH-H1 sequence, which cannot homo-dimerize on its own (29). The persistence of repressive function in non-nuclear proteins may suggest that there are moonlighting functions for more of these proteins.

### LisH domains are important for human disease

The human oncogene HsTBL1 is a transcriptional regulator and exchange factor involved in repression and activation (4, 22, 35, 36). Its dysregulation is implicated in the progression of multiple cancers (38, 50, 51). All human TBL1 isoforms (TBL1X, TBL1XR1, TBL1Y) contain a LisH domain, and the N-terminal region of TBL1X (residues 1-76, Fig. 3A) could replace TPL in our synthetic circuit (Figure 3B, dashed lines). As with TPL, TBL1-H1 repressed to a similar degree as the whole TBL1 N-terminus, and created a circuit that was more sensitive to auxin-induced degradation (Fig. 3B, solid lines).

**Figure 3:**
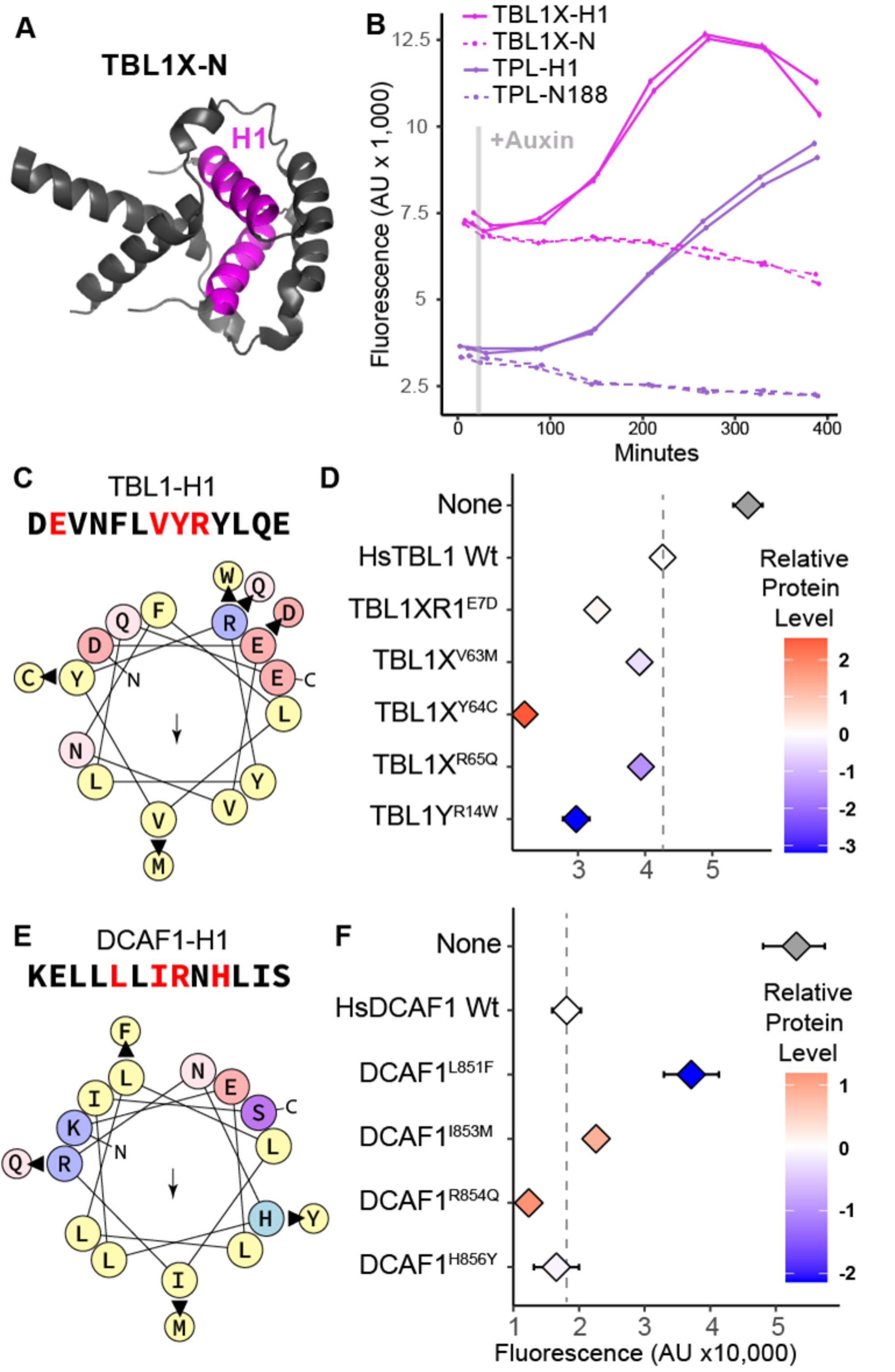
LisH domains are important for human disease. **A**. The dimerized protein structure of the N-terminal domain of Hs TBL1X, with H1 highlighted (pink, 2XTD). **B**. Time course flow cytometry of H1 and N-termini of HsTBL1X-IAA3 and AtTPL-IAA3 following auxin addition. Auxin (IAA-10µM) was added at the indicated time (gray bar, + Aux). HsTBL1X (pink) and AtTPL (purple), isolated H1 (solid line), N-terminal region (dotted line). Yeast strains are grown and measured after 400 minutes, with auxin added to some samples (solid lines) at the time indicated by the gray line. **C**. and **E**. Helical wheel depiction of HsTBL1X and HsDCAF1 H1 sequences colored by their physicochemical class (yellow – hydrophobic, red/blue – charged, pink/purple – polar uncharged, arrow indicates hydrophobic face) produced by HeliQuest (55). Arrows show the mutations found in these loci in the catalog of Somatic Mutations in Cancer (COSMIC) library (52), and where they occur. **D**,**F**. Effects on protein repressive function of these mutations in HsTBL1 and HsDCAF1 sequences are measured using flow cytometry. Each panel represents two independent time course flow cytometry experiments of the H1s indicated. For all cytometry, every point represents the average fluorescence of at least 10,000 individually measured yeast cells (a.u. - arbitrary units). Protein accumulation was measured by western blot and normalized to yeast PGK1. Levels of mutant protein constructs are shown relative to wild type H1s (HsTBL1-H1 and HsDCAF1-H1, respectively, dotted line) with each data point color coded from blue (low) to red (high) expression on a log2 scale.

The Catalogue of Somatic Mutations in Cancer (COSMIC) (52) has recorded five non-synonymous mutations in HsTBL1-H1 (pooled mutations from data for TBL1X, TBL1XR1, TBL1Y whose H1s are identical in sequence, Fig. 3C). We tested the functional impact of these mutations by engineering them into the HsTBL1-H1-substituted version of *At*ARC^Sc^. All tested variants increased HsTBL1 repressive function. HsTBL1-H1^Y64C^ was the strongest repressor and accumulated to the highest level, suggesting that this variant may potentiate oncogenesis by stabilizing HsTBL1. Both HsTBL1-H1^R65Q^ and HsTBL1-H1^R14W^ had lower protein abundance than wild-type HsTBL1-H1 yet higher repression rates. Together, these results imply that the level of repression by HsTBL1-H1 variants could be a useful, high throughput test for the disease-causing potential of a specific variant. Further investigation with many more variants, and associated clinical outcomes, is needed to follow-up on that prediction.

A number of LisH containing proteins in our phylogeny are components of E3 ubiquitin ligase complexes, one of which is a substrate receptor for Cullin RING ligase 4 (CRL4) and is named <u>D</u>DB1 (DNA damage-binding protein 1) and <u>C</u>UL4-associated factor <u>1</u> (DCAF1, (34). HsDCAF1 regulates diverse cell processes, and has been implicated in cancer (34) as well as subversion by HIV viral accessory proteins (33). The DCAF1 LisH has been implicated in both dimerization (21) and transcriptional repression, where it has been demonstrated to inhibit p53’s transcriptional activity through binding of hypoacetylated Histone 3 tails (53, 54). Following the same methodology as described for TBL1-H1, we engineered four known, non-synonymous cancer-associated mutations into the HsDCAF1-H1-substituted version of *At*ARC^Sc^ (Fig. 3E). HsDCAF1-H1 had a strong repressive function (Fig. 3F). Activity and fusion protein accumulation were affected minimally in HsDCAF1-H1^H856Y^ and HsDCAF1-H1^I853M^. However, HsDCAF1-H1^L851F^ dramatically reduced levels of the fusion protein and repressive function, while HsDCAF1-H1^R854Q^ was one of the strongest repressors that we tested and accumulated to a high level. More information is needed to determine whether any change in HsDCAF1-H1 measured in the *At*ARC^Sc^ is relevant in disease settings. The performance metrics of the variants tested for both HsTBL1-H1 and HsDCAF1-H1 could be applied to future design of synthetic repression domains, and screening for small molecule agonists/antagonists.

### LisH-H1s are effective synthetic repressor domains in plants

As LisH-H1s from distantly related species repressed transcription in yeast, we wondered whether they would work in other organisms as well. As a first test, we established a transient plant repression assay using essentially the same components as the *At*ARC^Sc^. The small number of modifications that we made to transfer the assay to plants included replacing yeast regulatory sequences for each gene with ones known to work in plants, switching to the well-characterized and highly sensitive DR5 auxin response reporter (56), and engineering mutations in IAA3 to prevent recruitment of endogenous TPL/TPR proteins (Fig. 4A). We found that, as in yeast, AtTPL-H1 was an effective repressor in the plant assays, and that similar trends were observed in the AtTPL-H1 variants tested in both systems (Fig. 4, Fig. 4 – Figure Supplement 7). Of particular interest for design of novel synthetic repressors, AtTPL-H1^F10A^ showed stronger repression than wild-type AtTPL-H1. We also tested sixteen of the most effective H1 repressors from our yeast assay, and successfully detected repression activity from nearly all of them (Fig 4). The few exceptions (e.g., Pf350509, DmCG5614) highlight the on-going challenge of standardizing parts across organisms. The overall success in transferring so many short repression domains from yeast to plants underscores the deep conservation of repression mechanisms across eukaryotes. It is also an excellent indicator that the LisH-H1 library characterized here—with its high tolerance for sequence diversity and range of repression strengths for tunability—can be a rich source of short, sequence-orthogonal repression domains for a diversity of engineering biology specifications.

**Figure 4.**
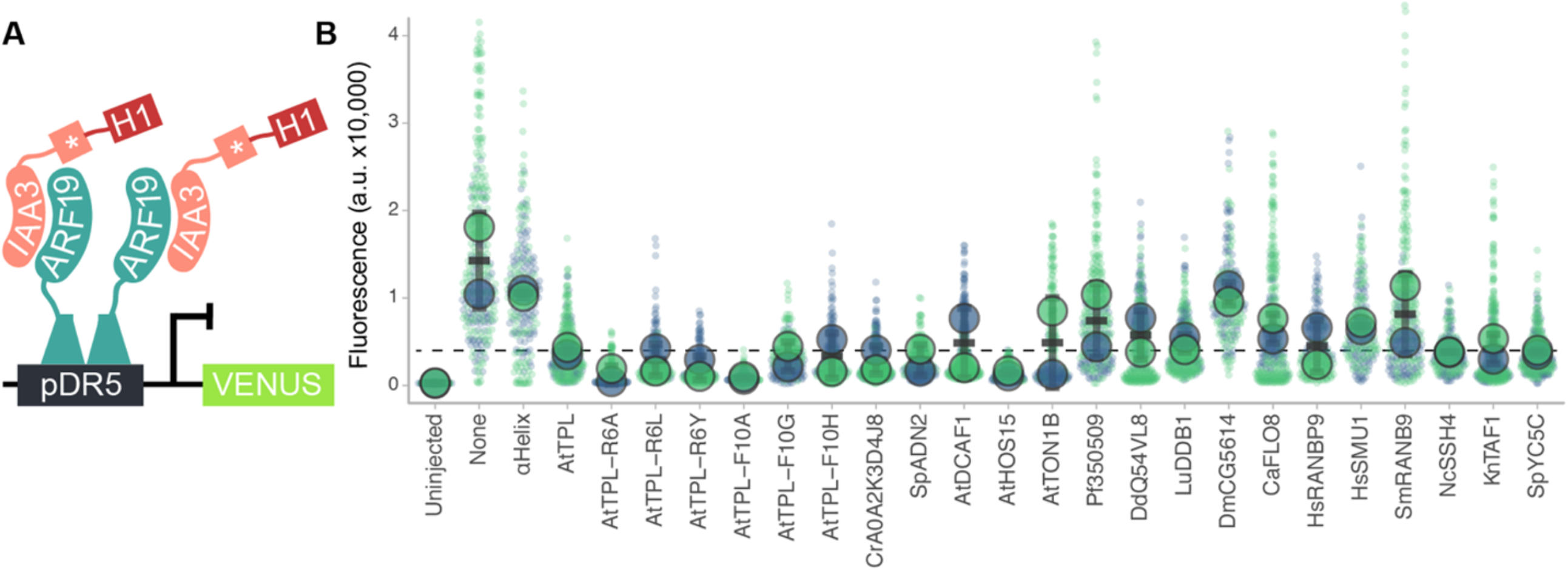
The H1 can act as a synthetic repressor domain *in planta*. **A**. Schematic of components injected in transient repression assays in *Nicotiana benthamiana*. pDR5:Venus reporter is based on a highly sensitive synthetic auxin promoter (57). Asterisk indicates EAR motif mutation. **B**. Repression activity of a range of H1 sequences are shown. Reporter activation was measured in four separate leaf injections (biological replicates) in two days of injection (large circles are pooled data from one day). Dashed line – AtTPL-H1 repression level.

## CONCLUSIONS

Synthetic signaling approaches aim to shed light on mechanisms of core cellular functions in natural systems, and to apply this knowledge to novel, beneficial interventions. The use of the *At*ARC^Sc^ in this study was successful for both objectives. First, we have isolated a single, short alpha helical domain within AtTPL that can function as an autonomous repression domain. Second, we have found that homologous domains found in thousands of proteins and diverse eukaryotes share this repression activity, and that repression was likely the function of the ancestral LisH-H1. Third, we were able to successfully apply knowledge of LisH-H1 function to the exploration of function of human disease-associated variants, as well as validate a number of these sequences as effective synthetic repression domains in plants. Future efforts should be directed at identifying the precise mechanism of transcriptional repression by AtTPL-H1, as well as other LisH-H1 sequences. These added insights would likely allow for a more directed approach to engineering LisH-H1 activity, as well as leading to a deeper understanding of what role LisH-H1 is playing in proteins that do not normally function in transcriptional regulation. A deepening of the connection between structure with function should also be useful for adapting the *At*ARC^Sc^ assays modified with human proteins for clinical applications, and optimizing the use of LisH-H1-based repression domains in synthetic circuits.

## METHODS

### Alignments and Phylogenetic Reconstruction

LisH domains were identified using UniProt (https://www.uniprot.org/), Pfam (http://pfam.xfam.org/family/PF08513.7) and SMART (http://smart.embl-heidelberg.de/) databases. LisH Helix 1 domains were aligned using Clustal Omega. Tree sequences were selected from the PFAM LisH clade PF08513 by performing an alignment of the representative proteome dataset with a 15% cutoff value (1235 sequences). The evolutionary history was inferred by using the Maximum Likelihood method and Le_Gascuel_2008 model (58). The tree with the highest log likelihood (−2711.40) was used. Initial tree(s) for the heuristic search were obtained automatically by applying Neighbor-Join and BioNJ algorithms to a matrix of pairwise distances estimated using the JTT model, and then selecting the topology with superior log likelihood value. A discrete Gamma distribution was used to model evolutionary rate differences among sites (5 categories (+*G*, parameter = 6.6850)). The percentage of replicate trees in which the associated taxa clustered together were calculated via bootstrap using 1000 replicates (59). This analysis involved 143 amino acid sequences. The cladogram was derived from this tree, only showing relationships among the 63 experimentally analyzed sequences. There was a total of 13 positions in the final dataset. Ancestral sequence reconstruction was done with an expanded tree using the same methods. Evolutionary analyses were conducted in MEGA X (42). Logo plots were created with an online tool ((60) https://weblogo.berkeley.edu/logo.cgi).

### Cloning

We used the VEGAS adapted method to create different forms of AtARC^Sc^ plasmids (61). We used a plasmid containing LisH-H1 fused to AtIAA13 and expressing URA3 as a backbone in LisH-H1 plasmid library construction. Plasmids were synthesized and confirmed with sequencing by Twist Bioscience (www.twistbioscience.com). All plasmids were transformed into reported haploid strain URA::[pRPS2-AtAFB2-ttCIT1, LEU2, pADH1-ARF19-ttADH1, pP3(2x)-UbiVenus-ttCYC1]. A standard lithium acetate protocol (62) was used for transformations of digested plasmids. All cultures were grown at 30°C with shaking at 220 rpm. For construction of plant vectors we used the MoClo toolkit (63) to design and clone plasmids containing our top 10 most repressive IAA13-H1s into vector pICH86966. Each H1 sequence is identical to yeast H1-HA-IAA3 constructs, except that IAA3 was mutated to prevent recruitment of endogenous TPL/TPR proteins. We transformed these into *A. tumefaciens* strain GV3101 via electroporation.

### Library Design

Phylogenetic library contains sequences selected from the Pfam LisH alignment PF08513 using the representative protein database (RP15, 1,235 sequences, http://pfam.xfam.org/family/PF08513). Ancestrally reconstructed sequences contain synthetic sequences predicted at nodes I-V using MEGAX node reconstruction software. HsTBL1-H1 and HsDCAF1-H1 mutational libraries contain somatic mutations found in human cancer cells within these helixes and were identified using COSMIC datasets ((52), https://cancer.sanger.ac.uk/cosmic). AtTPL site-saturation mutational libraries at residues AtTPL-H1 R6 and F10 contain synthetic sequences probing the function of these sites in helix 1. The alpha helix control sequence (EAAAK)_3_ was created based on well-studied synthetic alpha helix linkers (64).

### Flow Cytometry

Fluorescence measurements were taken using a Becton Dickinson (BD) special order cytometer with a 514-nm laser excitation fluorescence that is cut off at 525 nm prior to photomultiplier tube collection (BD, Franklin Lakes, NJ). Events were annotated, subset to singlet yeast using the FlowTime R package (https://github.com/wrightrc/flowTime). A total of 10,000 - 20,000 events above a 400,000 FSC-H threshold (to exclude debris) were collected for each sample and data exported as FCS 3.0 files for processing using the flowCore R software package and custom R scripts (65, 66). Data from at least two independent replicates were combined and plotted in R (https://ggplot2.tidyverse.org/).

### Yeast Methods

Standard yeast drop-out and yeast extract–peptone–dextrose plus adenine (YPAD) media were used, with care taken to use the same batch of synthetic dropout (SDO) media for related experiments. Haploid transformants were selected on appropriate prototrophy (SDO -Tryptophan, -Leucine). Yeast were grown at 30°C on selection plates for two days, and in SDO liquid media with 250rpm in a deep well 96-well plate format overnight for cytometry analysis (66). Liquid cultures were diluted 1:200 with fresh SDO the morning of cytometry analysis and measured after 5 hours of growth to a concentration of ∼200-500 events/μL.

### Western Blot

For yeast expressed proteins, yeast cultures grown to an OD600 of 1. Cells were harvested by centrifugation. Cells were lysed by vortexing for 5 min in the presence of 200 μl of 0.5-mm diameter acid washed glass beads and 200 μl SUMEB buffer (1% SDS, 8 M urea, 10 mM MOPS pH 6.8, 10 mM EDTA, 0.01% bromophenol blue, 1mM PMSF). Lysates were then incubated at 65°C for 10 min and cleared by centrifugation prior to electrophoresis and blotting. Antibodies: anti-HA-HRP (REF-12013819001, Clone 3F10, Roche/Millipore Sigma, St. Louis, MO), anti-PGK1 (ab113687, AbCam). Protein concentrations were quantified using ImageJ, with PGK1 protein measured in each strain to normalize protein concentrations across strains. To compare protein concentrations to AtTPL H1, these were then normalized to TPL concentration using this equation: ([X-H1]/[PGK1])/[AtTPL-H1]/[PGK1] where X is the H1 variant. AtTPL-H1 normalized protein concentrations were plotted on a Log2 scale. For tobacco expressed proteins, four leaf discs were pooled from one representative experiment and homogenized by bead-beating with two steel ball bearings for 1 minute. 200 μl of protein sample buffer was added to each sample, vortexed for 5 minutes and then boiled for 10 minutes and cleared by centrifugation prior to electrophoresis and blotting. Antibodies: anti-HA-HRP (REF-12013819001, Clone 3F10, Roche/Millipore Sigma, St. Louis, MO).

### Plant growth

For synthetic repression assays in tobacco, *Agrobacterium*-mediated transient transformation of *N. benthamiana* was performed as per (67). 5 ml cultures of *Agrobacterium* strains were grown overnight at 30°C shaking at 220 rpm, pelleted, and incubated in MMA media (10 mM MgCl2, 10 mM MES pH 5.6, 100 µM acetosyringone) for 3 hours at room temperature with rotation. Strain density was normalized to an OD600 of 1 for each strain in the final mixture of strains before injection into tobacco leaves. Leaves were removed, and eight different regions were excised using a hole punch, placed into a 96-well microtiter plate with 100 µl of water. Each leaf punch was scanned in a 4 × 4 grid for yellow and red fluorescence using a plate scanner (Tecan Spark, Tecan Trading AG, Switzerland). Fluorescence data was quantified and plotted in R (ggplots2).

## Acknowledgements

We thank members of the Nemhauser group including Cassandra Maranas, Eric Yang, and Dr. Sarah Guiziou for constructive discussions and comments on this manuscript. We thank Prof. Grant Brown and Prof. Maitreya Dunham for advice on yeast genetics and approaches.

**Supplement 1: the single plasmid ARC uses a hybrid integrated/unintegrated yeast auxin response circuit**. A. Flow cytometry on strains containing the ARC split into two plasmids with (light blue) or without (dark blue) an H1 repressor. In circuits with an unintegrated reporter, repression was observed, yet there was a wide peak width, limiting the resolution between the repressed and de-repressed response states. **B**. Integration of all components except the repressor led to tighter peak width distributions and increased the resolution of the repressed state when tested by fluorescence flow cytometry. **C**. Schematic of engineered versions of the H1-IAA3 repressor with single (1x) or double (2x) HA epitope tags, and Western blots with antibodies against HA and PGK1. A single HA epitope was sufficient for detection. **D**. Flow cytometry of epitope tagged H1-IAA constructs (1xHA – violet, 2xHA – wine, untagged repressors - light blue, no repressor - dark blue). **E**. Summary of fluorescence flow cytometry. For all flow cytometry each panel represents two independent time course flow cytometry experiments of the TPL helices indicated, all fused to IAA3, every plot represents the average fluorescence of at least 10,000 individually measured yeast cells (a.u. - arbitrary units).

**Supplement 2: Extended phylogeny for Figure 2**.The evolutionary history of LisH-H1 sequences was inferred by using the Maximum Likelihood method and Le Gascuel, 2008 model (58). The tree with the highest log likelihood (−2709.60) is shown. The percentage of trees in which the associated taxa clustered together is shown next to the branches. Initial tree(s) for the heuristic search were obtained automatically by applying Neighbor-Join and BioNJ algorithms to a matrix of pairwise distances estimated using the JTT model, and then selecting the topology with superior log likelihood value. A discrete Gamma distribution was used to model evolutionary rate differences among sites (5 categories (+G, parameter = 6.1034)). The tree is drawn to scale, with branch lengths measured in the number of substitutions per site. This analysis involved 143 amino acid sequences. There were a total of 13 positions in the final dataset. Evolutionary analyses were conducted in MEGA X (59). The percentage of replicate trees in which the associated taxa clustered together in the bootstrap test (1000 replicates) are shown next to the branches (42).

**Supplement 3: H1 sequence information**. Each column details information about **A**. each H1 sequence, including **B**. the gene names used for each sequence in figures, **C**. the function and **D**. localization of those proteins, **E**. A Uniprot-searchable name for each gene, **F**. the species each gene belongs to, **G**. other genes with an identical H1 sequence, **H**. and citations for annotated localization and function for each gene (“NONE” for those with uncharacterized function or localization). We also list all plasmids and oligos used.

**Supplement 4: Repression assay data visualizations**. The same cytometry data represented in Figure 2, ordered by A. how much repression we were able to detect, and B. how much protein accumulated. A, B. We have marked the fluorescence levels detected by the positive AtTPL-H1-HA-IAA3 control (dashed line) and negative IAA3 repression control (dotted line). Protein accumulation was measured by western blot and normalized to yeast PGK1. Levels of LisH-H1 proteins are shown relative to AtTPL-H1 with each data point color coded from blue (low) to red (high) expression on a log2 scale. For all cytometry, every point represents the average fluorescence of at least 10,000 individually measured yeast cells (a.u. - arbitrary units).

**Supplement 5: Clade logo plots**. Each panel represents the residues found in the H1s of proteins across the **A**. top 20 most repressive sequences, **B**. clade I, **C**. Clade II **D**. clade III, **E**. clade IV, **F**. and clade V. Taller columns represent more conserved residues. Letters appear longer the more commonly they are found at the specified residue. Letters at well-conserved residues are color coded by their physicochemical class. Logo plots were created with an online tool ((60) https://weblogo.berkeley.edu/logo.cgi).

**Supplement 6: Ancestral Sequence Reconstruction** A. The phylogeny from Supplement 1 was used to infer ancestral sequences at nodes marked in Supplement 1. A simplified cladogram shows these nodes (black dots) next to extant LisH-H1 sequences tested (black) and LisH-H1 sequences used to contextualize nodes (grey). Sequence clade I.I exists within clade representing an ancestral state closer to AtTPL-H1. Reconstructed sequences clade I and clade II are identical to Sc15387 and DdQ54VL8 respectively. B. Sequences were aligned and residues colored by their physicochemical class (RASMOL color scheme). The consensus sequence for LisH-H1 is aligned alongside the relative conservation rate of different residues along the helix. C. Flow cytometry and a modified *At*ARC^Sc^ depicted in Fig. 1E was used to quantify the relative repressive function of different H1s. For all cytometry, every point represents the average fluorescence of at least 10,000 individually measured yeast cells (a.u. - arbitrary units). We have marked the fluorescence levels detected by the positive AtTPL-H1-HA-IAA3 control (dashed line) and negative IAA3 repression control (dotted line). Protein accumulation was measured by western blot and normalized to yeast PGK1. Levels of construct expressions are shown relative to AtTPL-H1 with each data point color coded from blue (low) to red (high) expression on a log2 scale.

**Supplement 7: Protein expression in transient expression assays in tobacco**. Transient expression assays from Figure 4 were analyzed by Western blotting to determine the expression levels of H1-HA-IAA3 fusion proteins. Four leaf discs from were pooled from one representative experiment and analyzed by electrophoresis and blotting with antibodies against HA. Two exposures are provided to demonstrate detection of lowly expressed H1-HA-IAA3 variants. Protein normalization was calculated compared to the AtTPL-H1 wild type sample and is listed below each lane.

