## Supplementary figures and images for "A single helix repression domain is functional across eukaryotes"

### Supplement 1

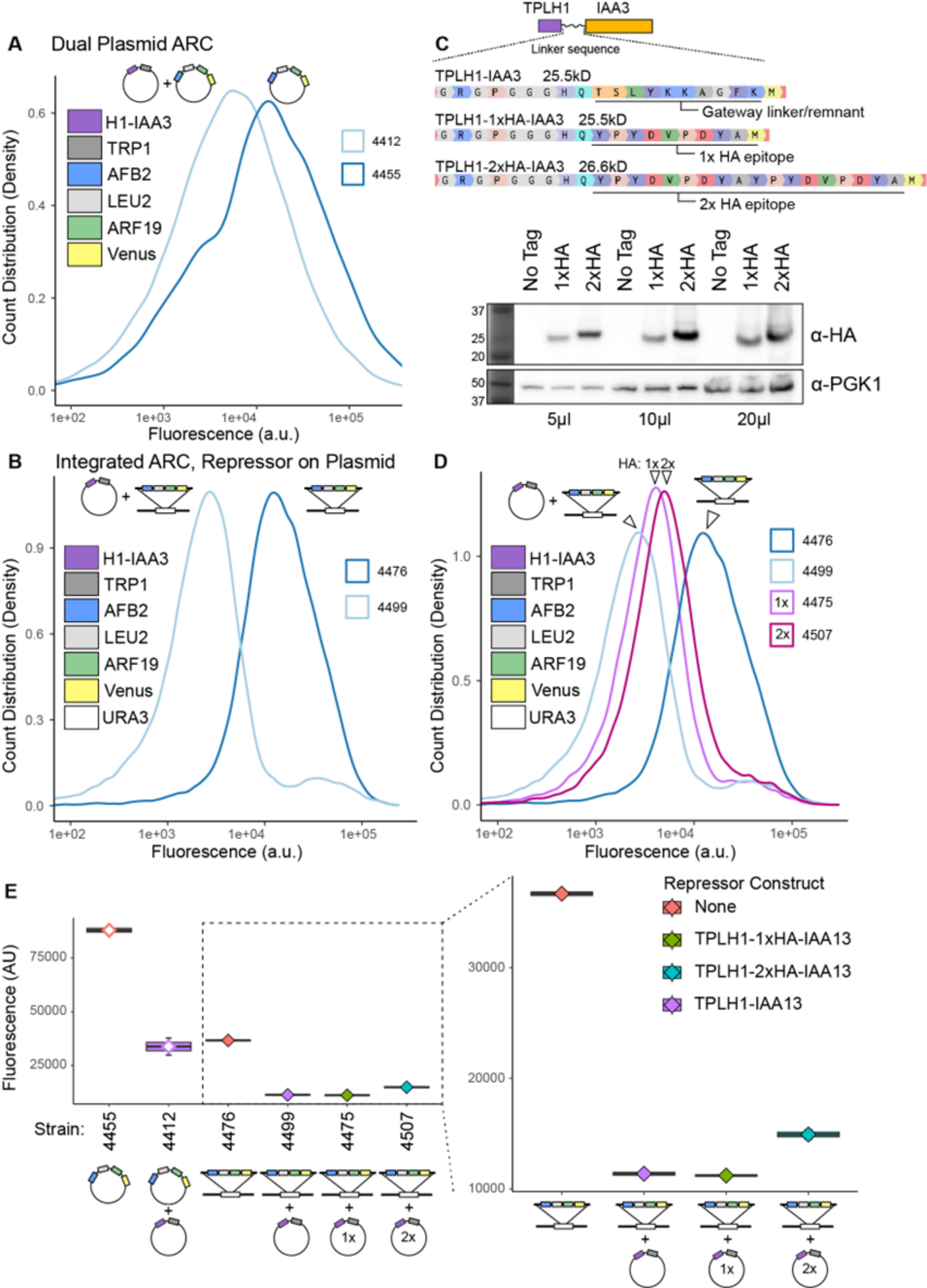

### Supplement 2

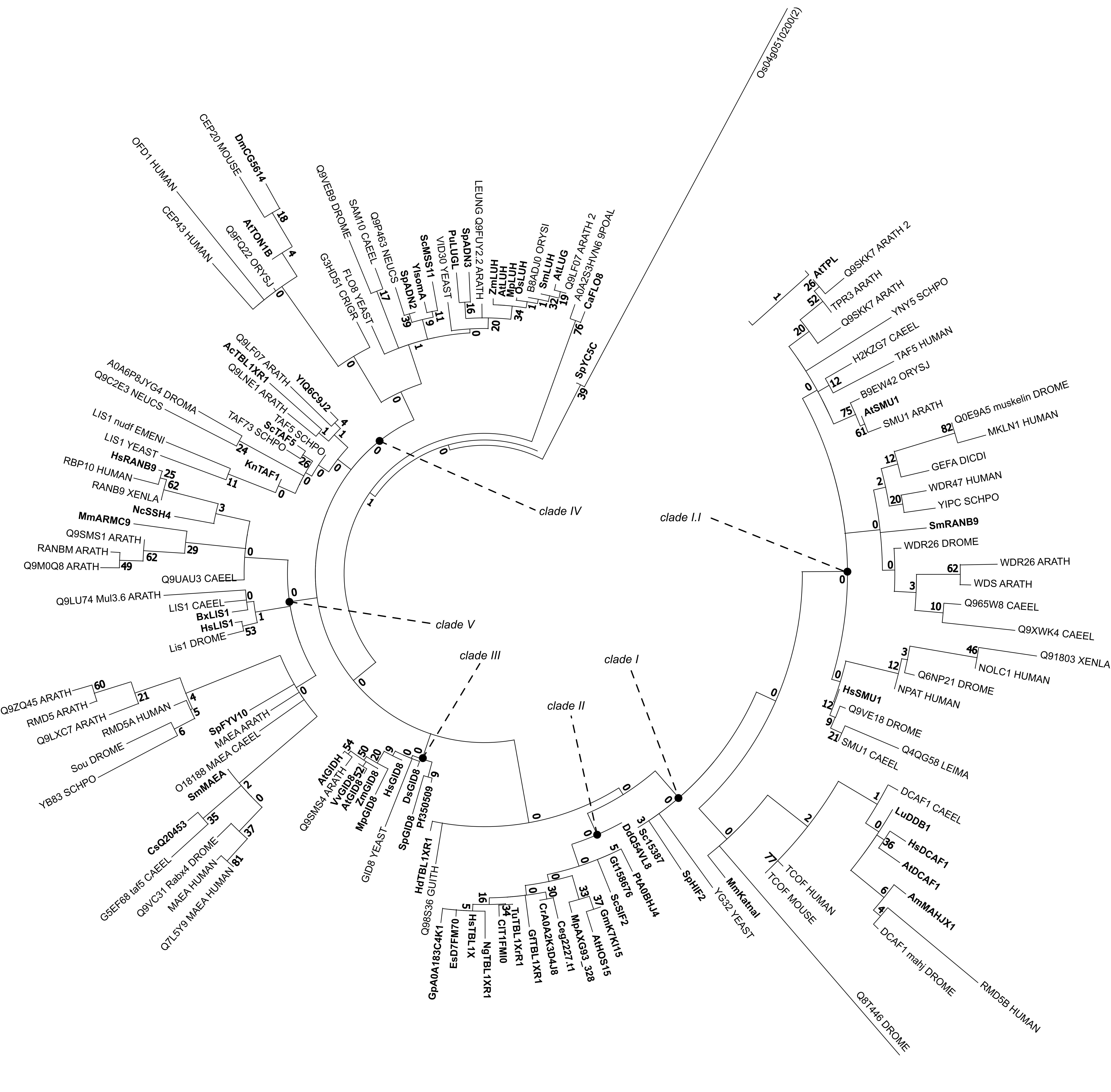

### Supplement 4

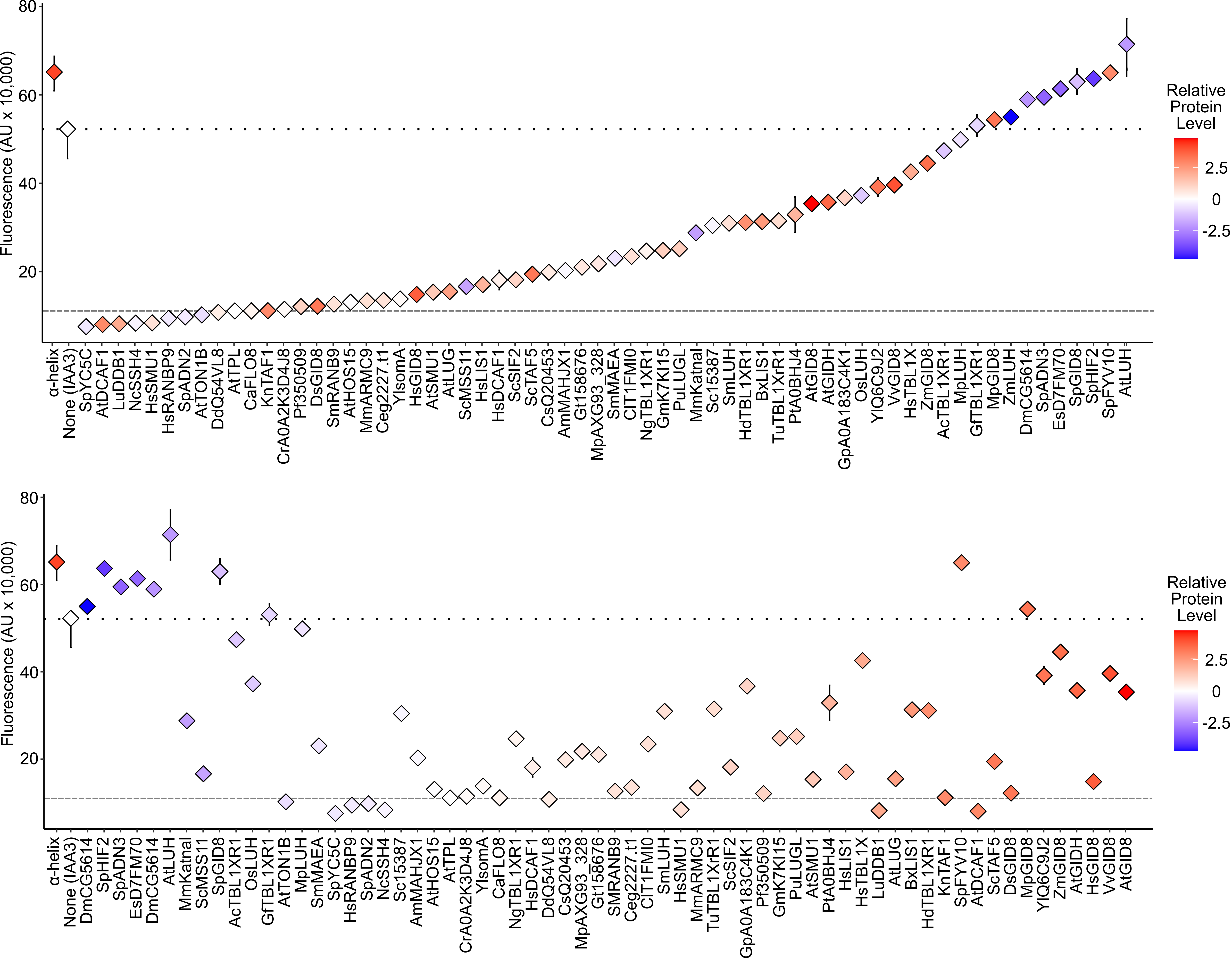

### Supplement 5

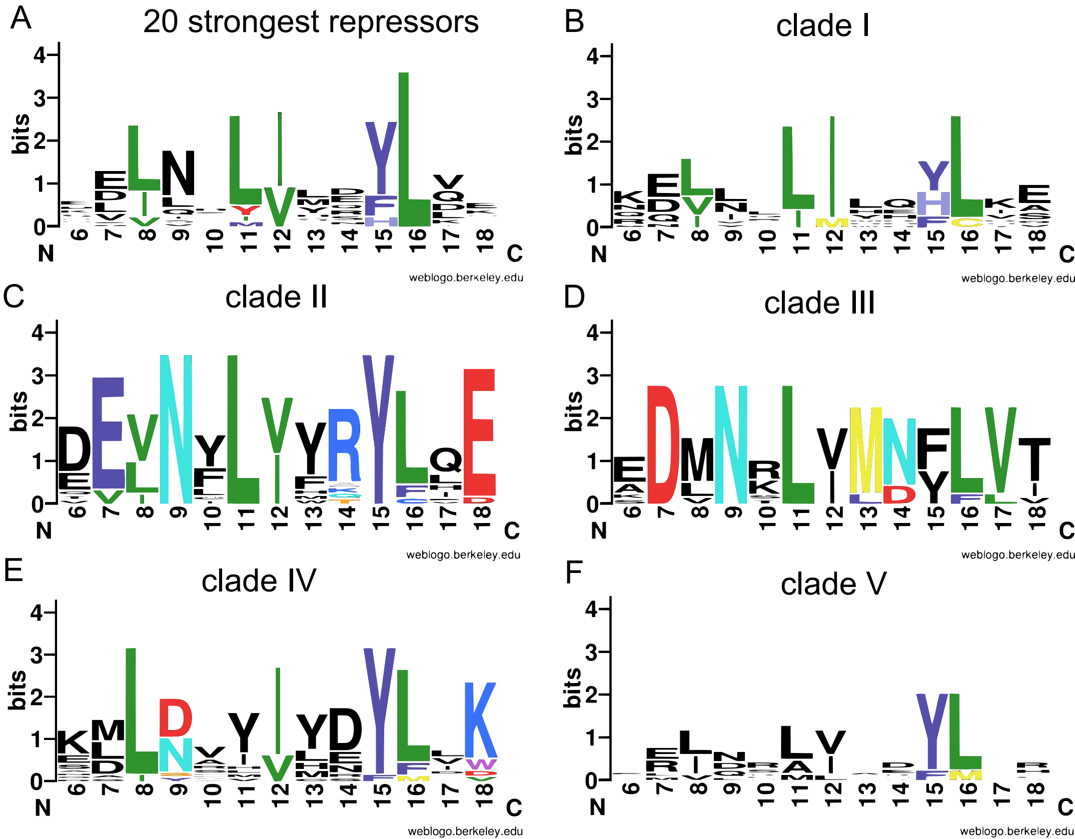

### Supplement 6

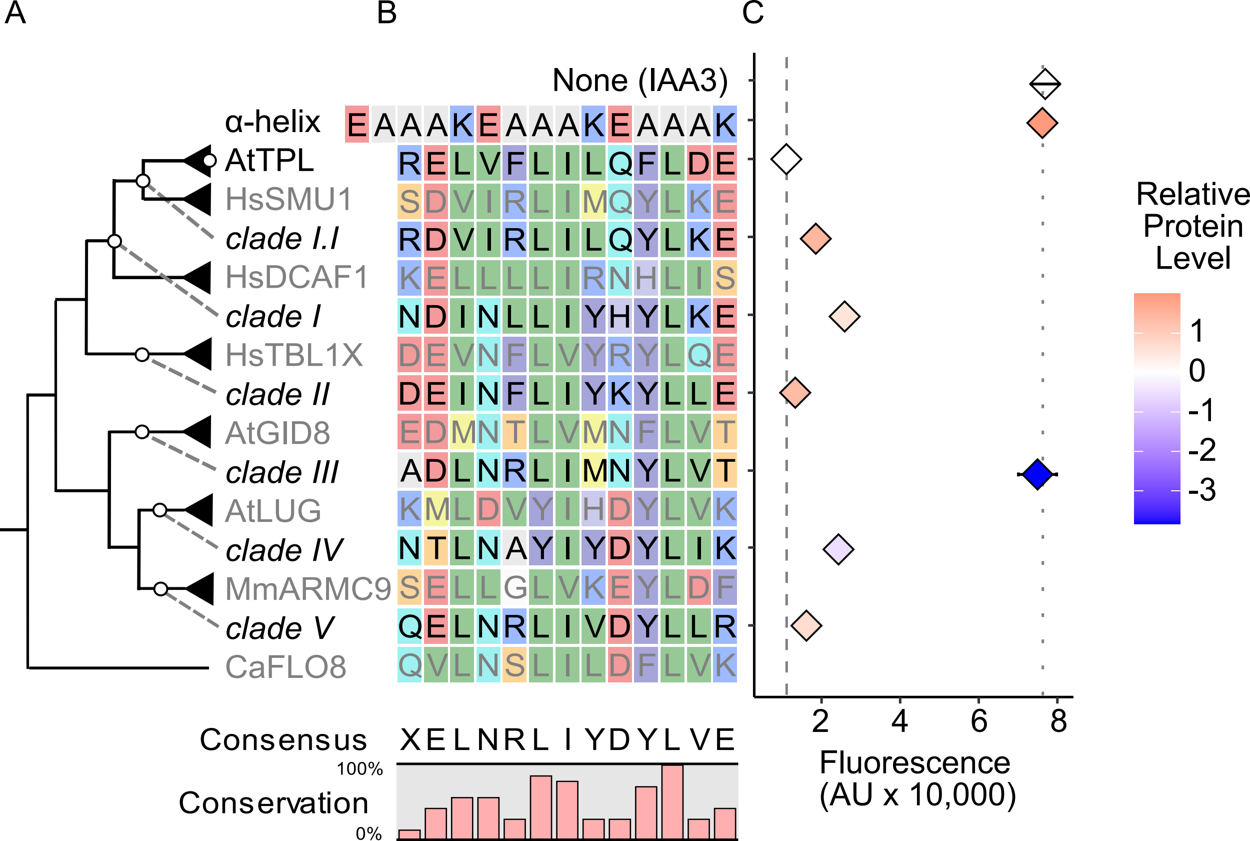

### Supplement 7

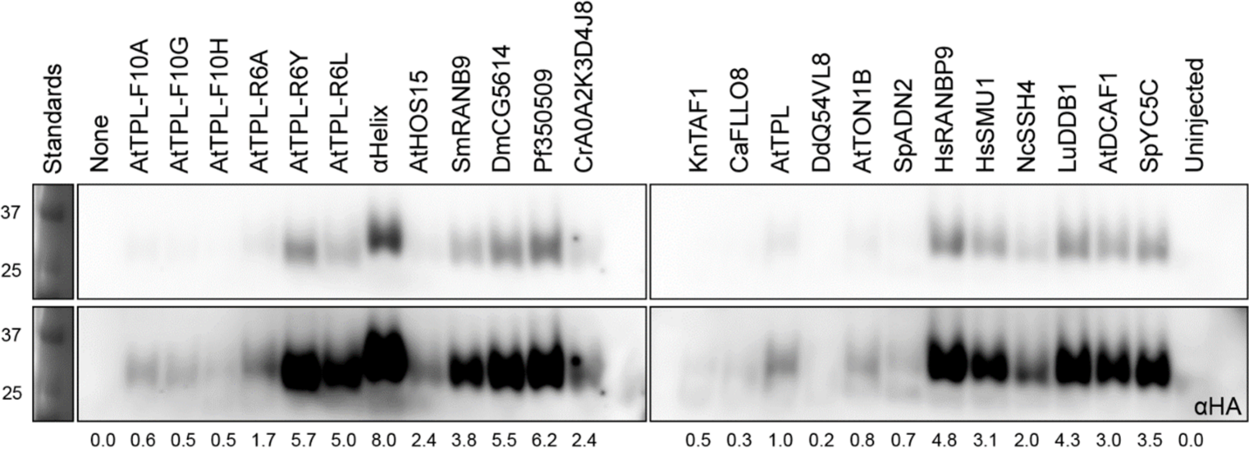
