## Supplement 3 for "A single helix repression domain is functional across eukaryotes"

| # | Sequence | Protein | function | localization | Uniprot Searchable | Species | Other genes | citation |
|---|---|---|---|---|---|---|---|---|
| 1 | RELVFLILQFLDE | AtTPL | Repressor | Nuclear | TPL_ARATH | Arabidopsis thaliana | TmTPL, VvTPR4, AoTPR2, CsTPL, CcTPR1, PaTPL, CmTPR2, AcTPR2, CnTPR3, TuTPR2, over 100 proteins | DOI: 10.1126/science.1151461 |
| 2 | RDVIKIMLQFCKE | AtSMU1 | Other | Nuclear | SMU1_ARATH | Arabidopsis thaliana | GwSMU1, HhSMU1, PpSMU1, HgSMU1, MdSMU1, BsSMU1X2, GaSMU1X1, GvSMU1, over 100 proteins | DOI: 10.1104/pp.109.141705 |
| 3 | QEINRLILEYLVV | **SmRANB9** | Unknown | Unknown | G4VEU8_SCHMA | *Schistosoma mansoni* | A0A183T2Q9, A0A068YDU5, H2KNK2, A0A183B251 , G4VEU8, A0A183VRA7, G4VEU8_SCHMA | NONE |
| 4 | SDVIRLIMQYLKE | HsSMU1 | Other | Nuclear | SMU1 | Homo sapiens (Human) | Q2TAY7 | DOI: 10.1016/j.yexcr.2005.02.017 |
| 5 | KELLLLIRNHLIS | HsDCAF1 | Other | Nuclear | Q9Y4B6 | Homo sapiens (Human) | PtDCAF1, CjDCAF1, NaDACF1, SaDCAF1, PcDACF1, and over 100 proteins | https://doi.org/10.1016/j.molcel.2012.09.004 |
| 6 | KELLLLIHEHLQA | AtDCAF1 | Other | Nuclear | DCAF1_ARATH | Arabidopsis thaliana | A0A178UYS1, Q9M086, DCAF1_ARATH | DOI: 10.1105/tpc.108.058891 |
| 7 | QQLYQLIYQHLIA | AmMAHJX1 | Unknown | Unknown | AgaP_AGAP012134 | Anopheles merus | AaMAHJX1, AcMAHJX1, AmMAHJ, AaMAHJX2, XP_320393.4, KFB51317.1 | NONE |
| 8 | KELLQLIHDHLVS | LuDDB1 | Unknown | Unknown | A0A1S3K6T0 | *Lingula unguis* | A0A1S3K7Q1, A0A1S3IUX2 | NONE |
| 9 | KNLLILILHYLTQ | MmKatnal | Other | Non-nuclear | KATL2_MOUSE | *Mus musculus* | F7DX12, F1M5A4, Q9D3R6, KATL2_MOUSE | DOI: 10.1371/journal.pgen.1007078 |
| 10 | NQVNYIIWRYLKE | SpHIF2 | Repressor | Nuclear | HIF2_SCHPO | Schizosaccharomyces pombe | none | DOI: 10.1016/s0960-9822(98)70304-5 |
| 11 | NDINLLIYHYLKE | Sc15387 | Unknown | Unknown | SteCoe_15387 | *Stentor coeruleus* | A0A1R2C3W4, A0A1R2CB06, A0A1R2BT11 | NONE |
| 12 | DEINFLIYKYLLE | DdQ54VL8 | Unknown | Unknown | DDB0206475_DICDI | *Dictyostelium discoideum* | DDB_G0280261, DICPUDRAFT_29050 | NONE |
| 13 | EELNYLIWRYCQE | ScSIF2 | Repressor | Nuclear | SIF2_YEAST | *Saccharomyces cerevisiae* | P38262, SIF2_YEAST | https://doi.org/10.1016/j.jmb.2005.06.025 |
| 14 | DELNYLIYRYLLE | **Gt158676** | Unknown | Unknown | GUITHDRAFT_158676 | *Guillardia theta* | BoCAD5229314.1 | NONE |
| 15 | EELNYLIYTYFLE | PtA0BHJ4 | Unknown | Unknown | A0BHJ4 | *Paramecium tetraurelia* | A0DLD6, A0BHJ4 | NONE |
| 16 | VELNFLVFRYLQE | AtHOS15 | Repressor | Nuclear | HOS15_ARATH | Arabidopsis thaliana | Q9FN19, A0A178UQT7, HOS15_ARATH, F2DCR6, A0A287I129, A0A287I0S4, A0A287I0S0 | DOI: 10.1104/pp.18.01156 |
| 17 | TELNYLVFRYLHE | GmK7KI15 | Unknown | Nuclear | K7KI15_GLYMA | Glycine max | A0A199W458, K7KI15, D8T4U4, D8RCD5 | NONE |
| 18 | QELNFLIFRYLHE | MpAXG93_328 | Unknown | Unknown | AXG93_328s1070 | *Marchantia polymorpha subsp. ruderalis* | A0A176VRL0 | NONE |
| 19 | DVVNYLIYRYLQE | CrA0A2K3D4J8 | Unknown | Unknown | A0A2K3D4J8_CHLRE | *Chlamydomonas reinhardtii* | TsFbxw7, CHLRE_12g527500v5, CiHXX76_004694, 10 proteins | NONE |
| 20 | DVINYLVMRYLQE | Ceg2227.t1 | Unknown | Unknown | CEUSTIGMA_g2227.t1 | *Chlamydomonas eustigma* | A0A250WW80 | NONE |
| 21 | DEVNCLIYAYFQD | GfTBL1XR1 | Unknown | Unknown | A0A1C7LUX2 | *Grifola frondosa* | KAH6916628.1 | NONE |
| 22 | DEVNYLVYRYLIE | ClT1FMI0 | Unknown | Unknown | T1FMI0_HELRO | *Helobdella robusta* | none | NONE |
| 23 | DEVNYLVYRYLQE | TuTBL1XrR1 | Unknown | Unknown | T1K4V1 | *Tetranychus urticae* | A0A2D4BVA7, C1FJI2, T1K4V1 | NONE |
| 24 | DEVNILVHRYLVE | NgTBL1XR1 | Unknown | Unknown | W7TZ27_9STRA | *Nannochloropsis gaditana* | NSK_003728 | NONE |
| 25 | DEVNLLVYRYLEE | GpA0A183C4K1 | Unknown | Unknown | GLOPA | *Globodera pallida* | MeCAD2177457.1, MgKAF7638710.1, MeCAD2168972.1 | NONE |
| 26 | EEVNFLVYRYLQE | EsD7FM70 | Unknown | Unknown | ECTSI | *Ectocarpus siliculosus* | A0A024G5P9, F0YLG7 | NONE |
| 27 | DEVNFLVYRYLQE | HsTBL1X | Activator/ Repressor | Nuclear | TBL1X_HUMAN | Homo sapiens (Human) | T0QLT6, D0NVN2, A7S5U0, A0A2B4S828, A0A1J1IKJ5, Q95RJ9, A0A1B0FD47, I0YVP9, H9JDI6, B7Q2X9, A0A1S3J4R8, A0A0L8H107, K1RDI6, V4A8I8, Q7Q371, A0A087UPF7, A0A2A4J0P6, A0A1S4EKS5, E9GI91, K7IPS0, A0A158NMP9, T1HP17, E0VC07, A0A2J7QTE6, A0A067R4E4, A0A226E5C7, A0A1W4XAC3, D6WDR4, A0A1D2N2Y2, T1JMU6, A0A2G8JKT3, A0A2G8LR25, C3ZEP1, O60907, S9XQS6, A2RUX0, A5WVU0, A0A1D5PXG9, A5PF33, F6S036, A0A226NGR2, Q9QXE7, G3V6G5, A0A1D5NXL1, Q9BQ87, Q9BZK7, A0A091KAK7, D3ZNF4, Q8BHJ5, M7B1G6, F6WUH3, W4XYV6, F6YCJ1 | DOI: 10.1210/jc.2016-2531 |
| 28 | EEVNLLIYQYLIE | HdTBL1XR1 | Unknown | Unknown | RAMVA_A0A1D1V3Z7 | *Hypsibius dujardini* | RAMVA_A0A1D1V3Z7 | NONE |
| 29 | KDLNKLIMNYLVI | **Pf350509** | Unknown | Unknown | BCR36DRAFT_350509 | *Piromyces finnis* | KAG4099912.1, OUM64527.1, ORY12276.1, | NONE |
| 30 | SDVNSLILDYLVI | SpGID8 | Unknown | Unknown | YDED_SCHPO | *Schizosaccharomyces pombe* | none | NONE |
| 31 | ADLNRLVMNYLVT | DsGID8 | Unknown | Unknown | B4IML9 | *Drosophila sechellia* | Q8SZN4 | NONE |

| | | | | | | | | |
|---|---|---|---|---|---|---|---|---|
| 32 | ADMNRLIMNYLVT | HsGID8 | Other | Nuclear | GID8_HUMAN | Homo sapiens (Human) | GID8_CHICK, A0A087TXU7, A0A1S3JEZ4, A0A226NJJ8, C3ZQT3, B7PS79, Q9VWS1, E9QHU7, Q6PC55, M7BF90, T1J9E3, Q9NWU2, Q9D7M1, Q6YDN8, F7CLX0, Q5ZKQ7, V8NIM7, F7DIR3, E7FGY2 | DOI: 10.7554/eLife.35528 |
| 33 | EDMNKLIMDFFLT | MpGID8 | Unknown | Unknown | MARPO_0028s0043 | Marchantia polymorpha | , AXG93_2931s1420 | NONE |
| 34 | EDMNRLVMNFLVT | ZmGID8 | Unknown | Unknown | A0A317YIJ7_MAIZE | Zea mays | , DKX38_017052, PR202_ga26473, Sadunf11G0026900, ZmGIH8, ONL95757.1, GFY91837.1, C3L33_15476, GFZ03098.1, C2845_PM01G08500, XP_003558414.1, 89 proteins | NONE |
| 35 | EDMNTLVMNFLVT | AtGID8 | Other | Other | GID8_ARATH | Arabidopsis thaliana | BRARA_K00274, CAF2133834.1, CAA7029683.1, Bca52824_086799, HID58_043154, DY000_02041394, BnaA03g60080D, XP_010418067.1, EsGIDH, CsGID8, CrGID8X1, 49 proteins | DOI: 10.1186/1471-2229-12-83 |
| 36 | EDMNKLVMNFLVT | VvGID8 | Unknown | Unknown | W1P668 | *Amborella trichopoda* | VvGIDH, KAG7019632.1, TEA_010235, E3N88_37174, FEM48_Zijuj01G0148200, Ahy_B03g063295, F2P56_009768, KAA0056956.1, KAE8716948.1, E3N88_41952, CcGID8, CcGIDH, CBI18750.3, 100 proteins | NONE |
| 37 | EDMNRLVMNFLVV | AtGIDH | Unknown | Unknown | O23690_ARATH | *Arabidopsis thaliana* | CAE5957163.1, AXX17_AT1G11330, KAG7591256.1, AlGID81X, KAG7653870.1, NP_001318979.1, EFH66107.1, KAG7645900.1, AAO63342.1, NP_001321853.1, CAD5312385.1, 19 proteins | NONE |
| 38 | ADLNSIVMDYLIV | YlQ6C9J2 | Unknown | Unknown | YALI0_D10791g | *Yarrowia lipolytica* | Q6C9J2 | NONE |
| 39 | KDLNLVVLRYLLD | AcTBL1XR1 | Unknown | Unknown | TBL1XR1_ANACO | *Ananas comosus* | CAD1841307.1, AcHOS15 | NONE |
| 40 | SDLNRIVLEYLNK | ScTAF5 | Activator | Nuclear | TAF5_YEAST | Saccharomyces cerevisiae | P38129, A0A1E3PKV2, W1Q9X6, A0A1D8PSS1, K0KTJ7, S6ENA4, A0A167EJ71, C4R4L4 | DOI: 10.1016/s0092-8674(00)81220-9 |
| 41 | EDLNRIVLSYLKK | KnTAF1 | Unknown | Unknown | A0A1Y1HYL3 | *Klebsormidium nitens* | A0A1Y1HYL3 | NONE |
| 42 | RLLSALICEYLDW | AtTON1B | Other | Other | TON1B_ARATH | Arabidopsis thaliana | CIPAW_02G177500, I3760_02G177300, JrTON1A, I3760_02G177300, CiTON1A, JmTON1A, Leryth_016134, CAD5325854.1, CmTON1AX1, AlTON1A, 44 proteins | doi: 10.1105/tpc.107.056812 |
| 43 | KLINQMIMEFLDW | DmCG5614 | Unknown | Unknown | Q9VF40 | *Drosophila melanogaster* | AT18024p | NONE |
| 44 | SLLNSYIYDYLIK | SpADN2 | Activator | Nuclear | ADN2_SCHPO | *Schizosaccharomyces pombe* | NONE | DOI: 10.1128/EC.00078-09 |
| 45 | ELLNAYIYDYLLK | YlsomA | Unknown | Unknown | YALI0_E03102g | *Yarrowia lipolytica* | FOA43_001062, HII12_001016, BKA90DRAFT_133770, B0I71DRAFT_126502, YALI2_E00070g, YALI0E03102p, BRETT_003633, CJU89_1027, 9 proteins | NONE |
| 46 | QLLYAHIYNYLIK | ScMSS11 | Activator | Nuclear | MSS11_YEAST | Saccharomyces cerevisiae | CAD6641633.1, AJS95620.1, Samss11p, YJM1443, YJM1389, JM1399, YJM320, AJS90831.1, AJS85596.1, SpMss11, AJS66828.1, AJS89087.1, AJS93898.1, AJS84716.1, 54 proteins | DOI: 10.1046/j.1365-2958.2003.03247.x |
| 47 | ESLDYYIYDYFVK | SpADN3 | Activator | Unknown | ADN3_SCHPO | *Schizosaccharomyces pombe* | ADN3_SCHPO | DOI: 10.1371/journal.pgen.1003104 |
| 48 | NALDFYIYNYMLK | PuLUGL | Unknown | Unknown | AXG93_146s1420 | Pyrus ussuriensis | MARPO_0023s0181 | NONE |

| | | | | | | | | |
|---|---|---|---|---|---|---|---|---|
| 49 | KMLDVYIYDYLVK | AtLUH | Activator/ Repressor | Nuclear | LUH_ARATH | Arabidopsis thaliana | A0A287T3L7, A0A287T3B8, A0A287T3E2, A0A287T3D2, A0A287T3C9, A0A287T3M2, A0A287T3B7, A0A199V044, Q10A93, K7K8T5, I1LA80, O48847, A0A178VQH2, A0A2K3P9I9, LUH_ARATH | DOI: 10.1105/tpc.19.00115, DOI: 10.1186/1471-2229-14-54 |
| 50 | KMLDVYIYDYFVK | ZmLUH | Unknown | Unknown | F2ELW0_HORVV | Zea mays | A0A287L2N7, A0A287L2N6, F2ELW0, A0A287L2M9, A0A287L2Q9, 75 proteins | NONE |
| 51 | KMLDVYIYDYLMK | MpLUH | Unknown | Unknown | MARPO_0033s0155 | Marchantia polymorpha | COLO4_34959, F8388_006400, DVH24_017861, RZB52406.1, RZB52407.1, PuLUGL, DVH24_029234, POE58772.1, QsLUG, C4D60_Mb07t20460, JHK85_023989, CAG1855999.1, CsLUG1X, 100 proteins | NONE |
| 52 | KMLDVYIYDYLLK | OsLUH | Unknown | Unknown | Q0JBT9 | *Oryza sativa subsp. japonica* | TuLUG, MQM01692.1, ZmLEGLC5167_041388, CY35_06G035600, BDL97_06G036400, C5167_015243, MQL71672.1, OIW03074.1, XP_019457374.1, LaLUGL, 100 proteins | NONE |
| 53 | KMLDVYIHDYLVK | AtLUG | Activator/ Repressor | Nuclear | LUG_ARATH | Arabidopsis thaliana | B9RPC5_RICCO, A0A2J6KP98, A0A178UVW2, A0A251SRG4, A0A251UB06, A0A0R0FPP1, A0A0R0JDA7, A0A2K3MNX1, D8SAR0, D8QW58, S8CL12, I1M9M2, S8DZ67, Q9FUY2 | DOI: 10.1073/pnas.230352397 |
| 54 | KMLDVYIHDYLTK | **SmLUH** | Unknown | Unknown | SELMO_94317 | *Selaginella moellendorffii* | SmLUGX1, X2, X3, X4 | NONE |
| 55 | EELNSALAEYLQR | BxLIS1 | Unknown | Unknown | A0A1I7STR7 | *Bursaphelenchus xylophilus* | , CAD5215296.1 | NONE |
| 56 | DELNRAIADYLRS | HsLIS1 | Other | Non-nuclear | LIS1_HUMAN | Homo sapiens (Human) | Q803D2, LIS1_BOVIN, Q7T394, Q9PTRZ, A0A091JVU8, P43034, P63005, P63004, F6PQP7, V8P437, Q6NZH4 | https://doi.org/10.1091/mbc.e12-03-0210 |
| 57 | QRVDRLIIDYLLR | SmMAEA | Unknown | Unknown | A0A0N5AP93 | *Syphacia muris* | VDD86692.1 | NONE |
| 58 | ERIDSLIYGYLRR | CsQ20453 | Unknown | Unknown | Q20453_CAEEL | *Caenorhabditis elegans* | H2L2B2, EGT35366.1, NP_001256138.1, NP_001256137.1, NP_505524.2, NP_872162.2, NP_001023935.1 | NONE |
| 59 | VRLNRLVADYMMA | SpFYV10 | Unknown | Nuclear | FYV10_SCHPO | Schizosaccharomyces pombe | NONE | DOI: 10.1038/nbt1222 |
| 60 | SELLGLVKEYLDF | MmARMC9 | Other | Other | ARMC9 MOUSE | Mus musculus | MXQ79359.1, MdARMC9, MbARMC9, KAF4014044.1, XP_037705437.1, XP_037705438.1, XP_045054028.1, XP_045054029.1, XP_022423649.1, NaARMC9X1, PtARMC9X1, DlARMC9X2, 100 proteins | DOI: 10.1038/s41588-018-0054-7 |
| 61 | ELIQQLVLQFLQH | NcSSH4 | Unknown | Unknown | Q7S7G6 | Neurospora crassa | GQX73_g5821, CIB48_g11750, PODANS_2_950, KAH6631079.1, PaRBP10, INS49_012262, SMAC_08383, NEUTE1DRAFT_57335, NcRANB, 43 proteins | NONE |
| 62 | TMIQKMVSSYLVH | HsRANBP9 | Other | Nuclear | RANB9_HUMAN | Homo sapiens (Human) | F1LVV3, P69566, M7BR83, Q96S59, A0A091KE14, F7D8G8, A0A226NKE1 | DOI: 10.1083/jcb.200801133 |
| 63 | QVLNSLILDFLVK | CaFLO8 | Activator | Nuclear | FLO8_CANAL | *Candida albicans* | XP_002421174.1, W5Q_05038 , AAQ03244.1, MGS_04938, W5O_04937, MG1_04938, MEU_04910, L150_04850, MEY_04890, MGO_04880, MEW_04821, MEM_04918, 29 proteins | DOI: 10.1091/mbc.e05-06-0502 |
| 64 | EFLNELISSFLLN | SpYC5C | Unknown | Unknown | YC5C_SCHPO | *Schizosaccharomyces pombe* | O94712 | NONE |

| YEAST PLASMIDS | | Corresponding H1 sequences |
|---|---|---|
| 3455 | c1-1_TPL_ARATH | RELVFLILQFLDE |
| 3454 | c9-2_DCAF1_ARATH | KELLLLIHEHLQA |
| 3453 | c17-3_LUH_ARATH | KMLDVYIYDYLVK |
| 3452 | c17-6_LUG_ARATH | KMLDVYIHDYLVK |
| 3451 | c14-4_HOS15_ARATH | VELNFLVFRYLQE |
| 3450 | c4-3_GID8_ARATH | EDMNTLVMNFLVT |
| 3448 | c4-8_O23690_ARATH | EDMNRLVMNFLVV |
| 3447 | c1-2_SMU1_ARATH_3 | RDVIKIMLQFCKE |
| 3446 | c20-1_TON1B_ARATH | RLLSALICEYLDW |
| 3445 | c15-9_TBL1X_HUMAN | DEVNFLVYRYLQE |
| 3444 | c18-2_LIS1_HUMAN | DELNRAIADYLRS |
| 3443 | c4-2_GID8_HUMAN | ADMNRLIMNYLVT |
| 3442 | c22-1_RANB9_HUMAN | TMIQKMVSSYLVH |
| 3441 | c10-1_KATL2_MOUSE | KNLLILILHYLTQ |
| 3440 | c1-3_SMU1_HUMAN | SDVIRLIMQYLKE |
| 3439 | c4-1_Dmel_DROME | ADLNRLVMNYLVT |
| 3438 | c19-1_MAEA_SYPMU | QRVDRLIIDYLLR |
| 3437 | c9-3_AgaP_AGAP012134 | QQLYQLIYQHLIA |
| 3436 | c15-5_HELRO | DEVNYLVYRYLIE |
| 3435 | c18-1_Lis1_BURXY | EELNSALAEYLQR |
| 3434 | c9-1_DDB1_LINUN | KELLQLIHDHLVS |
| 3433 | c15-6_T1K4V1 | DEVNYLVYRYLQE |
| 3432 | c11-1_RAMVA | EEVNLLIYQYLIE |
| 3431 | c20-2_Q9VF40 | KLINQMIMEFLDW |
| 3430 | c15-7_GLOPA | DEVNLLVYRYLEE |
| 3429 | c3-1_BCR36DRAFT_350509 | KDLNKLIMNYLVI |
| 3428 | c23-1_YC5C_SCHPO | EFLNELISSFLLN |
| 3427 | c14-1_SIF2_YEAST | EELNYLIWRYCQE |
| 3426 | c5-1_FLO8_CANAL | QVLNSLILDFLVK |
| 3425 | c8-1_TAF5_YEAST | SDLNRIVLEYLNK |
| 3424 | c16-3_MSS11_YEAST | QLLYAHIYNYLIK |
| 3423 | c17-1_ADN3_SCHPO | ESLDYYIYDYFVK |
| 3422 | c12-1_HIF2_SCHPO | NQVNYIIWRYLKE |
| 3421 | c16-2_YALI0_E03102g | ELLNAYIYDYLLK |
| 3420 | c15-3_Tbl1xr1_RHVIN | DEVNCLIYAYFQD |
| 3419 | c16-1_ADN2_SCHPO | SLLNSYIYDYLIK |
| 3418 | c6-1_YALI0_D10791g | ADLNSIVMDYLIV |
| 3417 | c19-3_FYV10_SCHPO | VRLNRLVADYMMA |
| 3416 | c3-2_YDED_SCHPO | SDVNSLILDYLVI |
| 3415 | c21-1_SSH4_NUECR | ELIQQLVLQFLQH |
| 3414 | c4-6_A8HQD2_CHLRE | EDMNRLVMNFLVT |
| 3413 | c15-1_A0A2K3D4J8_CHLRE | DVVNYLIYRYLQE |
| 3412 | c4-5_AMTR | EDMNKLVMNFLVT |
| 3411 | c4-4_MARPO_0028s0043 | EDMNKLIMDFFLT |
| 3410 | c17-5_MARPO_0033s0155 | KMLDVYIYDYLMK |
| 3409 | c7-1_TBL1XR1_ANACO | KDLNLVVLRYLLD |
| 3408 | c17-7_SELMO_94317 | KMLDVYIHDYLTK |
| 3407 | c14-2_GUITH_158676 | DELNYLIYRYLLE |
| 3406 | c24-1_Q0JBT9 | KMLDVYIYDYLLK |
| 3405 | c17-4_F2ELW0_HORVV | KMLDVYIYDYFVK |
| 3404 | c15-4_tbl1xr1_NAGAD | DEVNILVHRYLVE |

| | | |
|---|---|---|
| 3403 | c17-2.2_AXG93_328s1070 | QELNFLIFRYLHE |
| 3402 | c15-2_CHEUS | DVINYLVMRYLQE |
| 3401 | c14-5_K7KI15_GLYMA | TELNYLVFRYLHE |
| 3400 | c17-2.1_AXG93_146s1420 | NALDFYIYNYMLK |
| 3399 | c8-2_TAF1_KLENI | EDLNRIVLSYLKK |
| 3398 | c15-8_ECTSI | EEVNFLVYRYLQE |
| 3397 | c2-1_RANB9_SCHMA | QEINRLILEYLVV |
| 3396 | c19-2_CELE_CAEEL | ERIDSLIYGYLRR |
| 3395 | c14-3_A0BHJ4 | EELNYLIYTYFLE |
| 3394 | c13-1_DDB0206475_DICDI | DEINFLIYKYLLE |
| 3393 | c12-2_SteCoe_15387 | NDINLLIYHYLKE |
| 3392 | TBL1XR1_E7D | DDVNFLVYRYLQE |
| 3391 | TBL1X_F61V | DEVNVLVYRYLQE |
| 3390 | TBL1X_V63M | DEVNFLMYRYLQE |
| 3389 | TBL1X_Y64C | DEVNFLVCRYLQE |
| 3388 | TBL1X_R65Q | DEVNFLVYQYLQE |
| 3387 | TBL1X_R14W | DEVNFLVYWYLQE |
| 3386 | HsDCAF1_wt | KELLLLIRNHLIS |
| 3385 | HsDCAF1_L851F | KELLFLIRNHLIS |
| 3384 | HsDCAF1_I853M | KELLLLMRNHLIS |
| 3383 | HsDCAF1_R854Q | KELLLLIQNHLIS |
| 3382 | HsDCAF1_H856Y | KELLLLIRNYLIS |
| 3381 | TPLH1_R6 | RELVFLILQFLDE |
| 3380 | TPLH1_R6H | HELVFLILQFLDE |
| 3379 | TPLH1_R6K | KELVFLILQFLDE |
| 3378 | TPLH1_R6D | DELVFLILQFLDE |
| 3377 | TPLH1_R6E | EELVFLILQFLDE |
| 3376 | TPLH1_R6S | SELVFLILQFLDE |
| 3375 | TPLH1_R6T | TELVFLILQFLDE |
| 3374 | TPLH1_R6N | NELVFLILQFLDE |
| 3373 | TPLH1_R6Q | QELVFLILQFLDE |
| 3372 | TPLH1_R6C | CELVFLILQFLDE |
| 3371 | TPLH1_R6G | GELVFLILQFLDE |
| 3370 | TPLH1_R6P | PELVFLILQFLDE |
| 3369 | TPLH1_R6A | AELVFLILQFLDE |
| 3368 | TPLH1_R6I | IELVFLILQFLDE |
| 3367 | TPLH1_R6L | LELVFLILQFLDE |
| 3366 | TPLH1_R6M | MELVFLILQFLDE |
| 3365 | TPLH1_R6F | FELVFLILQFLDE |
| 3364 | TPLH1_R6W | WELVFLILQFLDE |
| 3363 | TPLH1_R6Y | YELVFLILQFLDE |
| 3362 | TPLH1_R6V | VELVFLILQFLDE |
| 3356 | ARMC9_Mm | SELLGLVKEYLDF |
| 3355 | alphahelix | EAAAKEAAAKEAAAK |
| 3353 | TPLH1_F10H | RELVHLILQFLDE |
| 3352 | TPLH1_F10D | RELVDLILQFLDE |
| 3351 | TPLH1_F10E | RELVELILQFLDE |
| 3350 | TPLH1_F10T | RELVTLILQFLDE |
| 3349 | TPLH1_F10N | RELVNLILQFLDE |
| 3348 | TPLH1_F10Q | RELVQLILQFLDE |
| 3347 | TPLH1_F10C | RELVCLILQFLDE |

| | | |
|---|---|---|
| 3346 | TPLH1_F10G | RELVGLILQFLDE |
| 3345 | TPLH1_F10A | RELVALILQFLDE |
| 3344 | TPLH1_F10I | RELVILILQFLDE |
| 3343 | TPLH1_F10L | RELVLLILQFLDE |
| 3341 | TPLH1_F10W | RELVWLILQFLDE |
| 3340 | TPLH1_F10V | RELVVLILQFLDE |
| 3339 | TPL_2xH1 | RELVFLILQFLDEGGGGSGGGGSRELVF LILQFLDE |
| 3338 | TPL_3xH1 | RELVFLILQFLDEGGGGSGGGGSRELVF LILQFLDEGGGGSGGGGSRELVFLILQF LDE |
| 3337 | TPL_F10A | RELVALILQFLDE |
| 3336 | TPL_F10R | RELVRLILQFLDE |
| 3335 | TPLH1_REDE>AAAA | AALVFLILQFLAA |
| 3334 | TPL_4xH1 | RELVFLILQFLDEGGGGSGGGGSRELVF LILQFLDEGGGGSGGGGSRELVFLILQF LDEGGGGSGGGGSRELVFLILQFLDE |
| 4714 | Clade I.I | RDVIRLILQYLKE |
| 4713 | Clade III | ADLNRLIMNYLVT |
| 4712 | Clade IV | NTLNAYIYDYLIK |
| 4711 | Clade V | QELNRLIVDYLLR |
| **AGRO PLASMIDS** | | **Corresponding H1 sequences** |
| 3548 | p35SLong_5U:[H1-AtTPL-F10A]-HA-IAA3-3U+Ter-AtuNos | RELVALILQFLDE |
| 3547 | p35SLong_5U:[H1-AtTPL-F10G]-HA-IAA3-3U+Ter-AtuNos | RELVGLILQFLDE |
| 3546 | p35SLong_5U:[H1-AtTPL-F10H]-HA-IAA3-3U+Ter-AtuNos | RELVHLILQFLDE |
| 3545 | p35SLong_5U:[H1-AtTPL-R6A]-HA-IAA3-3U+Ter-AtuNos | AELVFLILQFLDE |
| 3544 | p35SLong_5U:[H1-AtTPL-R6Y]-HA-IAA3-3U+Ter-AtuNos | YELVFLILQFLDE |
| 3543 | p35SLong_5U:[H1-AtTPL-R6L]-HA-IAA3-3U+Ter-AtuNos | LELVFLILQFLDE |
| 3542 | p35SLong_5U:[H1-alphaH]-HA-IAA3-3U+Ter-AtuNos | EAAAKEAAAKEAAAK |
| 3541 | p35SLong_5U:[H1-AtHOS15]-HA-IAA3-3U+Ter-AtuNos | VELNFLVFRYLQE |
| 3540 | p35SLong_5U:[H1-RANB9-schma]-HA-IAA3-3U+Ter-AtuNos | QEINRLILEYLVV |
| 3539 | p35SLong_5U:[H1-DmDMEI]-HA-IAA3-3U+Ter-AtuNos | ADLNRLVMNYLVT |

| | | |
|---|---|---|
| 3538 | p35SLong_5U:[H1-BCR36DRAFT]-HA-IAA3-3U+Ter-AtuNos | KDLNKLIMNYLVI |
| 3537 | p35SLong_5U:[H1-A0A2K3D4J8]-HA-IAA3-3U+Ter-AtuNos | DVVNYLIYRYLQE |
| 3536 | p35SLong_5U:[H1-TAF1]-HA-IAA3-3U+Ter-AtuNos | EDLNRIVLSYLKK |
| 3535 | p35SLong_5U:[H1-FLO8]-HA-IAA3-3U+Ter-AtuNos | QVLNSLILDFLVK |
| 3534 | p35SLong_5U:[H1-AtTPL]-HA-IAA3-3U+Ter-AtuNos | RELVFLILQFLDE |
| 3533 | p35SLong_5U:[H1-DDB0206475]-HA-IAA3-3U+Ter-AtuNos | DEINFLIYKYLLE |
| 3532 | p35SLong_5U:[H1-AtTON1B]-HA-IAA3-3U+Ter-AtuNos | RLLSALICEYLDW |
| 3531 | p35SLong_5U:[H1-ADN2]-HA-IAA3-3U+Ter-AtuNos | SLLNSYIYDYLIK |
| 3530 | p35SLong_5U:[H1-HsRANBP9]-HA-IAA3-3U+Ter-AtuNos | TMIQKMVSSYLVH |
| 3529 | p35SLong_5U:[H1-HsSMU1]-HA-IAA3-3U+Ter-AtuNos | SDVIRLIMQYLKE |
| 3528 | p35SLong_5U:[H1-SSH4]-HA-IAA3-3U+Ter-AtuNos | ELIQQLVLQFLQH |
| 3527 | p35SLong_5U:[H1-DDB1]-HA-IAA3-3U+Ter-AtuNos | KELLQLIHDHLVS |
| 3526 | p35SLong_5U:[H1-AtDCAF1]-HA-IAA3-3U+Ter-AtuNos | KELLLLIHEHLQA |
| 3525 | p35SLong_5U:[H1-YC5C]-HA-IAA3-3U+Ter-AtuNos | EFLNELISSFLLN |
| **Name** | **OLIGO SEQUENCE** | **Purpose** |
| TPLH1_R6A_F | TTCTCTTAGTgctGAGCTCGTTTTCTTG | AtTPL-H1 Alanine mutations |
| TPLH1_R6A_R | GACATCATTTTGTGATGGATC | AtTPL-H1 Alanine mutations |
| TPLH1_E7A_F | TCTTAGTAGAgctCTCGTTTTCTTGATC | AtTPL-H1 Alanine mutations |
| TPLH1_E7A_R | GAAGACATCATTTTGTGATG | AtTPL-H1 Alanine mutations |
| TPLH1_R6AE7A_F | TTCTCTTAGTgctgctCTCGTTTTCTTGATCTTACAG | AtTPL-H1 Alanine mutations |
| TPLH1_R6AE7A_R | GACATCATTTTGTGATGGATC | AtTPL-H1 Alanine mutations |
| TPLH1_F10A_F | AGAGCTCGTTgctTTGATCTTACAGTTTC | AtTPL-H1 Alanine mutations |

| | | |
|---|---|---|
| TPLH1_F10A_R | CTACTAAGAGAAGACATCATTTTG | AtTPL-H1 Alanine mutations |
| TPLH1_Q14A_F | CTTGATCTTAgctTTTCTCGATGAAGGGAGAGG | AtTPL-H1 Alanine mutations |
| TPLH1_Q14A_R | AAAACGAGCTCTCTACTAAG | AtTPL-H1 Alanine mutations |
| TPLH1_D17A_F | ACAGTTTCTCgctGAAGGGAGAG | AtTPL-H1 Alanine mutations |
| TPLH1_D17A_R | AAGATCAAGAAAACGAGCTC | AtTPL-H1 Alanine mutations |
| TPLH1_E18A_F | GTTTCTCGATgctGGGAGAGGAC | AtTPL-H1 Alanine mutations |
| TPLH1_E18A_R | TGTAAGATCAAGAAAACGAG | AtTPL-H1 Alanine mutations |
| TPLH1_D17AE18A_F | ACAGTTTCTCgctgctGGGAGAGGAC | AtTPL-H1 Alanine mutations |
| TPLH1_D17AE18A_R | AAGATCAAGAAAACGAGC | AtTPL-H1 Alanine mutations |
| TPLN_1xHA_F | gccggattatgcgTCTTCTCTTAGTAGAGAGC | AtTPL-H1 HA tag addition |
| TPLN_1xHA_R | acatcatacggataCATCATTTTGTGATGGATC | AtTPL-H1 HA tag addition |
| TPLN_2xHA_F | tatccgtatgatgtgccggattatgcgTCTTCTCTTAGTAGAGAGC | AtTPL-H1 HA tag addition |
| TPLN_2xHA_R | cgcataatccggcacatcatacggataCATCATTTTGTGATGGATC | AtTPL-H1 HA tag addition |
| 3132_F10X_R | CTACTAAGAGAAGACATCATTTTG | AtTPL-H1 F10 mutations |
| 3132_F10E_F | AGAGCTCGTTgaaTTGATCTTACAGTTTC | AtTPL-H1 F10 mutations |
| 3132_F10Q_F | AGAGCTCGTTcaaTTGATCTTACAGTTTC | AtTPL-H1 F10 mutations |
| 3132_F10R_F | AGAGCTCGTTagaTTGATCTTACAGTTTC | AtTPL-H1 F10 mutations |
